# Remarkable sex-specific differences at Single-Cell Resolution in Neonatal Hyperoxic Lung Injury

**DOI:** 10.1101/2022.08.19.504541

**Authors:** A Cantu, M Cantu, X Dong, C Leek, E Sajti, K Lingappan

## Abstract

Exposure to supraphysiological concentrations of oxygen (hyperoxia) predisposes to bronchopulmonary dysplasia (BPD), which is characterized by abnormal alveolarization and pulmonary vascular development, in preterm neonates. Neonatal hyperoxia exposure is used to recapitulate the phenotype of human BPD in murine models. Male sex is considered an independent predictor for the development of BPD, but the main mechanisms underlying sexually dimorphic outcomes are unknown. Our objective was to investigate sex-specific and cell-type specific transcriptional changes that drive injury in the neonatal lung exposed to hyperoxia at single-cell resolution and delineate the changes in cell-cell communication networks in the developing lung. We used single cell RNA sequencing (scRNAseq) to generate transcriptional profiles of >35000 cells isolated from the lungs of neonatal male and female C57BL/6 mice exposed to 95% FiO2 between PND1-5 (saccular stage of lung development) or normoxia and euthanized at PND7 (alveolar stage of lung development). ScRNAseq identified 22 cell clusters with distinct populations of endothelial, epithelial, mesenchymal, and immune cells. Our data identified that the distal lung vascular endothelium (composed of aerocytes and general capillary endothelial cells) is exquisitely sensitive to hyperoxia exposure with the emergence of an intermediate capillary endothelial population with both aCaP and gCaP markers. We also identified a myeloid derived suppressor cell population from the lung neutrophils. Sexual dimorphism was evident in all lung cell subpopulations but was striking among the lung immune cells. Finally, we identified that the specific intercellular communication networks and the ligand-receptor pairs that are impacted by neonatal hyperoxia exposure.

## Introduction

The male disadvantage for neonatal mortality and major morbidities in preterm neonates is well known (1–4). Sex-chromosome-based, hormonal, imprinting, or epigenetic mechanisms may modulate the male susceptibility or the female resilience (5). Susceptibility to injury and the subsequent repair and recovery may differ between the sexes due to the sexual dimorphism at baseline or pathways activated or inhibited in response to the injurious stimuli.

Poor lung health in premature neonates is a major factor underlying an inferior quality of life, economic costs related to health care and risk of developing adult-onset chronic lung diseases (6, 7). Respiratory morbidity including the development of bronchopulmonary dysplasia (BPD) is common in preterm neonates with long term impact. Even in the post-surfactant era, extremely premature male neonates (born between 24 and 26 weeks of gestation) displayed a significantly increased risk of respiratory complications (8). The incidence of respiratory distress syndrome (RDS), BPD, and moderate to severe BPD was significantly increased in males after adjusting for multiple confounding factors, particularly in the 750-999 gms birth weight group (9). Similarly, the need for respiratory support, respiratory medications and home oxygen use was higher in males (10, 11). Male sex was an independent risk for tracheostomy among preterm neonates and in infants who received a tracheostomy, an independent predictor of mortality (12). Hospital readmissions and outpatient visits due to respiratory issues in the 1-4 years and 5-9 years age groups were also increased in male premature neonates (13). Male sex is thus an independent predictor for the development of BPD and the subsequent morbidities related to lung disease in premature infants, but the underlying mechanisms behind these sexually dimorphic outcomes are unknown.

Animal models for human BPD use several approaches to replicate the major findings of alveolar simplification and abnormal vascular remodeling in the murine lung including postnatal hyperoxia exposure in the term mouse (14). Exposure to hyperoxia and the resulting oxidative stress to the developing lung contributes to the pathophysiology of this disease. The murine lung is in the saccular stage of lung development from birth to post-natal day (PND4-5), which is equivalent to 26-36 weeks in human preterm neonates (15). Most preterm neonates are exposed to varying degrees of hyperoxia in the neonatal intensive care unit during this period, with the sicker neonates receiving the highest concentrations of oxygen support (16).

Our lab and others have reported sex-specific differences in alveolar and vascular development in neonatal mice using this model (17–19). Furthermore, we have highlighted differences involving epigenetic mechanisms, micro-RNA mediated effects and the role sex-chromosome and gonadal hormones in this model all of which show striking sex-specific differences (20–23). However, the sex-specific differences in individual lung cell sub populations at single-cell resolution at the saccular stage of lung development in the setting of neonatal hyperoxia exposure are not known. In this investigation, we tested the hypothesis that there are marked sex-specific differences in different lung cell subpopulations in the developing murine lung during the saccular stage of lung development upon exposure to hyperoxia. Significantly, we show that distal lung vascular endothelium composed of the *aCaP* (aerocytes) and the *gCaP* (general capillary) endothelial cells, is exquisitely sensitive to hyperoxia exposure with the emergence of an intermediate capillary endothelial population with both *aCaP* and *gCaP* endothelial cells and the identification of a myeloid derived suppressor cell population from the lung neutrophils. Sexual dimorphism was evident in all lung subpopulations but was striking among the lung immune cells. Finally, we identified that the specific intercellular communication networks and the ligand-receptor pairs that are impacted by neonatal hyperoxia exposure.

## RESULTS

### Single cell RNA sequencing of healthy and hyperoxia exposed lungs in postnatal day 7

To generate a comprehensive study of the sex specific transcriptional changes after hyperoxia exposure in the early lungs, one day old male and female mice (day of birth) were exposed to room air (21% Oxygen) or hyperoxia (95% Oxygen) until day 5. Mice were allowed to recover in room air until day 7 when lungs were collected and processed for single cell RNA sequencing (**Figure 1A**). A total of 35,934 cell transcriptomes were profiled with cell clustering showing capture overlap between room air and hyperoxia cells in both male and female lungs (**Figure 1B**). *Epcam+* cells were identified as epithelial, *Cldn5*+ as endothelial, *Ptprc*+ as immune and *Col1a2+* as mesenchymal cells (24) (**Figure 1C-D**). Additionally, 22 clusters were identified by comparing their expression profile to published literature (25–30) (**Figure 1C, E**). The number of cells sequenced in each group and the highly expressed genes in each cluster are included in **Supplemental Table 1.**

**Figure 1.**
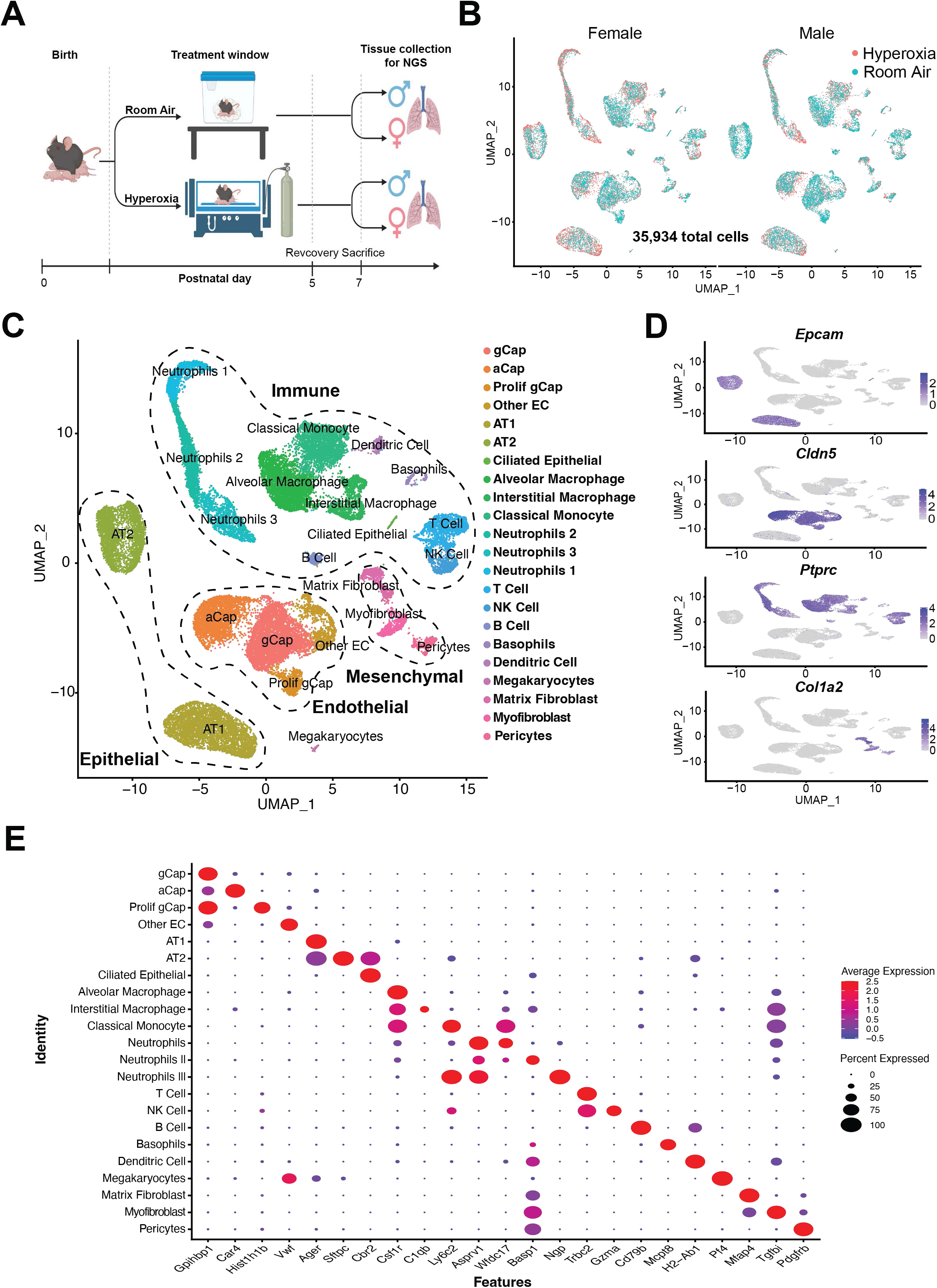
Capturing the effects of hyperoxia in the lung at single cell resolution. **A)** Experimental design and timeline. Mouse pups (C57BL6) from multiple litters were pooled before being randomly and equally redistributed to two groups, one group exposed to room air (21% O2) and the other group exposed to hyperoxia (95% O2), within 12 h of birth for 5 days, and euthanized on PND7 **B)** UMAP of sequenced cells labeled by experimental condition (room air in teal or hyperoxia in red) and split by sex (male or female). **C)** UMAP overview of cell clusters identified based on gene expression. Dotted groups encompass large cell groups of distinct populations separated by endothelial, epithelial, immune and mesenchymal. **D)** UMAP plot of expression levels of cell markers for epithelial (*Epcam*), endothelial (*Cldn5*), immune (*Ptprc*) and mesenchymal (*Col1a2*) cell populations. **E)** Dot plot showing expression levels (dot color) and percent of cells expressing each gene (dot size) of previously validated marker genes for each cluster identified.

### Endothelial expression changes in response to hyperoxia

Sub clustering of 9,394 endothelial cells identified seven distinct populations (**Figure 2A**). Alveolar specific general capillaries (gCap) and aerocytes or alveolar capillaries (aCap) make up over 75% of the cells captured. gCap cells express the transporter for lipoprotein lipase *Gpihbp1,* while actively proliferating gCap cells express the cell cycle genes like *Hist1h1b* (Figure 1E) (25). aCap cells were identified by their expression of carbonic anhydrase 4 or *Car4* (Figure 1E) (25). All large vessels were marked by expression of von Willebrand factor (*vWF*, Figure 1E) (31) with specific expression of *Gja4* in arterial, *Nr2f2* in venous and *Prox1* (25) in lymphatic cells. Interestingly, a second population of aCap cells, that we called reactive aCap, were characterized by the upregulation of *Inhba* (24) and greatly expand after hyperoxia exposure and represent more than 50% of the total lung endothelial cells (**Figure 2B-C**).

**Figure 2.**
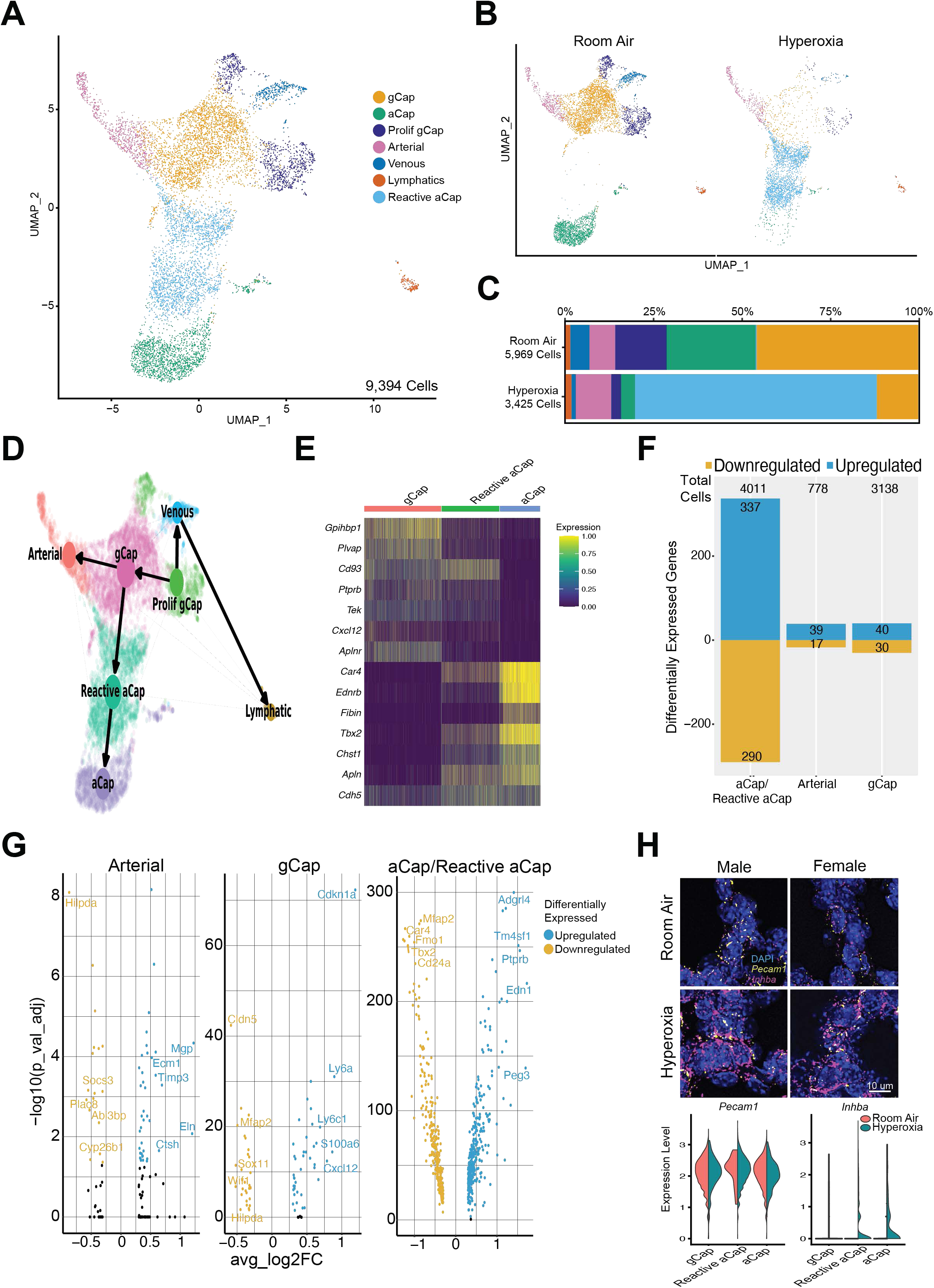
Gene Expression Changes in Endothelial Cells in Response to Hyperoxia. **A)** UMAP of sequenced lung endothelial cells identified seven distinct clusters. **B)** UMAP of lung endothelial cells showing the seven distinct clusters in room air and hyperoxia. **C)** Changes in the relative contribution of different lung endothelial cell sub-populations in room air and hyperoxia. **D)** Trajectory analysis of lung endothelial cells showing reactive aCaPs emerging from gCaPs and leading to aCaPs. **E)** Gene expression similarities and differences between aCaP, reactive aCaPs and gCaPs in the distal lung microvasculature. **F)** Number of cells sequenced and the number of up- and down-regulated genes in aCaP/reactive aCaP, gCaP and arterial endothelial cells upon exposure of the neonatal lung to hyperoxia. **G)** Volcano plots showing the differentially expressed genes in arterial, gCaP and aCaP/reactive aCaP endothelial cells. **H)** In situ hybridization showing the increased expression of *Inhba* in lung endothelial cells in response to hyperoxia and violin plots showing increased expression of *Inhba* in aCaP and reactive aCaP endothelial cells.

To understand the expression dynamics between the reactive aCaps and the rest of the endothelial cell population we employed RNA Velocity (32) and partition-based graph abstraction. RNA Velocity results were consistent with previously published data showing that venous cells give rise to lymphatic endothelium (33) as well as proliferative gCap giving rise to mature gCap cells (25). Interestingly, we observed the gCap cluster specifying into reactive aCap cells that then further connected into aCap cells suggesting that reactive aCap is an intermediate less differentiated cell state that occurs in response to injury (**Figure 2D**). Gene expression in aCaPs, gCaPs and reactive aCaPs is shown in a heat map in **Figure 2E**. Reactive aCaPs have an expression profile that is similar in some respects to aCaPs (high *Car4* and high *Apln*) and some to gCaPs (high *CD93*).

Differential expression shows the largest number of genes changing in aCap/reactive aCaP cells (**Figure 2F-G**). We observed 337 genes upregulated and 290 downregulated in hyperoxia. *Inhba* expression was increased in reactive aCaPs upon exposure to hyperoxia and this was validated using single molecule fluorescent in situ hybridization (smFISH), with higher expression of *Inhba* among *Pecam* positive cells in the hyperoxia exposed lung compared to room air controls (**Figure 2H**). Our finding was reported previously by Hurskainen *et al.*(*24*) in a newborn hyperoxia model after 14 days of hyperoxia exposure (85% FiO2). Another upregulated gene target that was identified and validated in the reactive aCaP population was paternally expressed gene (*Peg) 3*. *Peg 3* activates autophagy in endothelial cells and attenuates angiogenesis through thromspondin-1 (34–37). We show increased expression of *Peg3* in aCaP cells (labelled by endothelial receptor b; *Ednrb*) in the hyperoxia exposed lung (**Supp Figure 1A**).

The second endothelial population with most gene expression changes were the gCap cells, where hyperoxia induced the downregulation of 30 genes and upregulation of 40 genes (**Figure 2F**). As the stem cell-like population, gCap cells are involved during lung repair (25) and their total number decrease lungs exposed to hyperoxia (**Figure 2C**). Interestingly, we observed the upregulation of *Ly6a or Sca1 (Stem cell antigen-1)* in gCap cells (*Aplnr+*) after hyperoxia (**Supp Figure 1B**). *Sca1* is used to mark endothelial progenitor cells (38) and is expressed in the large vessel and capillary endothelium in the lung (39). *Sca 1+* endothelial gCaP cells may contribute to endothelial repair after injury (40, 41).

In arterial cells, the endothelial homeostasis regulator *Socs3 (Suppressor of cytokine signaling 3)* (42) is downregulated in hyperoxia while pro-angiogenic factor *Ecm1 (Extracellular matrix protein 1)* (*43*) is upregulated (**Figure 2G**).

Biological pathways that were differentially regulated in aCaPs/reactive aCaPs and gCaP cells in the distal capillary endothelium were identified. The upregulated pathways that were common to male and female aCaPs/reactive aCaPs included oxidative phosphorylation, ECM-receptor interaction, cytokine-mediated receptor signaling and mTORC1 signaling (**Supp Figure 1C**). The switch from glycolysis to oxidative phosphorylation in endothelial cells under pathological conditions, is well described(44–46). Downregulated pathways included regulation of ERK1 and ERK2 and TGF-beta receptor signaling pathway. ERK signaling plays a crucial role in angiogenesis and in maintaining endothelial quiescence(47–49). The gCaPs were positively enriched for pathways related to p53 signaling, mesenchymal transition, while regulation of cellular proliferation was downregulated (**Supp Figure 1D**). Differentially expressed genes in all endothelial cell sub-populations are listed in **Supplemental Table 2.**

### Sex specific response to hyperoxia in endothelial cells

Next, we separated our cells by sex to explore the gene expression changes of the lung endothelium in response to hyperoxia injury (**Figure 3A**). Differentially expressed genes up- and down-regulated in aCaps/reactive aCaPs, gCaPs and arterial endothelial cells in male and female lungs are shown in **Figure 3B**. Interestingly, the number of sex-specific genes was greater than the shared genes in these endothelial cell subpopulations. Volcano plots and biological pathways in male and female aCaPs/reactive aCaps and gCaPs are shown in **Figures 3C and 3D** respectively. *Klf 4* (Kruppel-like factor 4) was downregulated in male aCaPs/reactive aCaps, *Klf 4* (*50*) modulates angiogenesis through the regulation of the Notch pathway. *Klf4* limits activation of Notch 1. *Ddah1 (*Dimethylarginine Dimethylaminohydrolase 1) was upregulated in the female acaP/reactive aCaPs. Loss of *Ddah 1* impairs while overexpression(51, 52) enhances angiogenesis. Sex-specific biological pathways included cellular senescence and artery morphogenesis downregulated and nitric oxide biosynthesis upregulated in female, and mitochondrial electron transport, response to oxidative stress and cholesterol homeostasis upregulated in males (Figure 3D).

**Figure 3.**
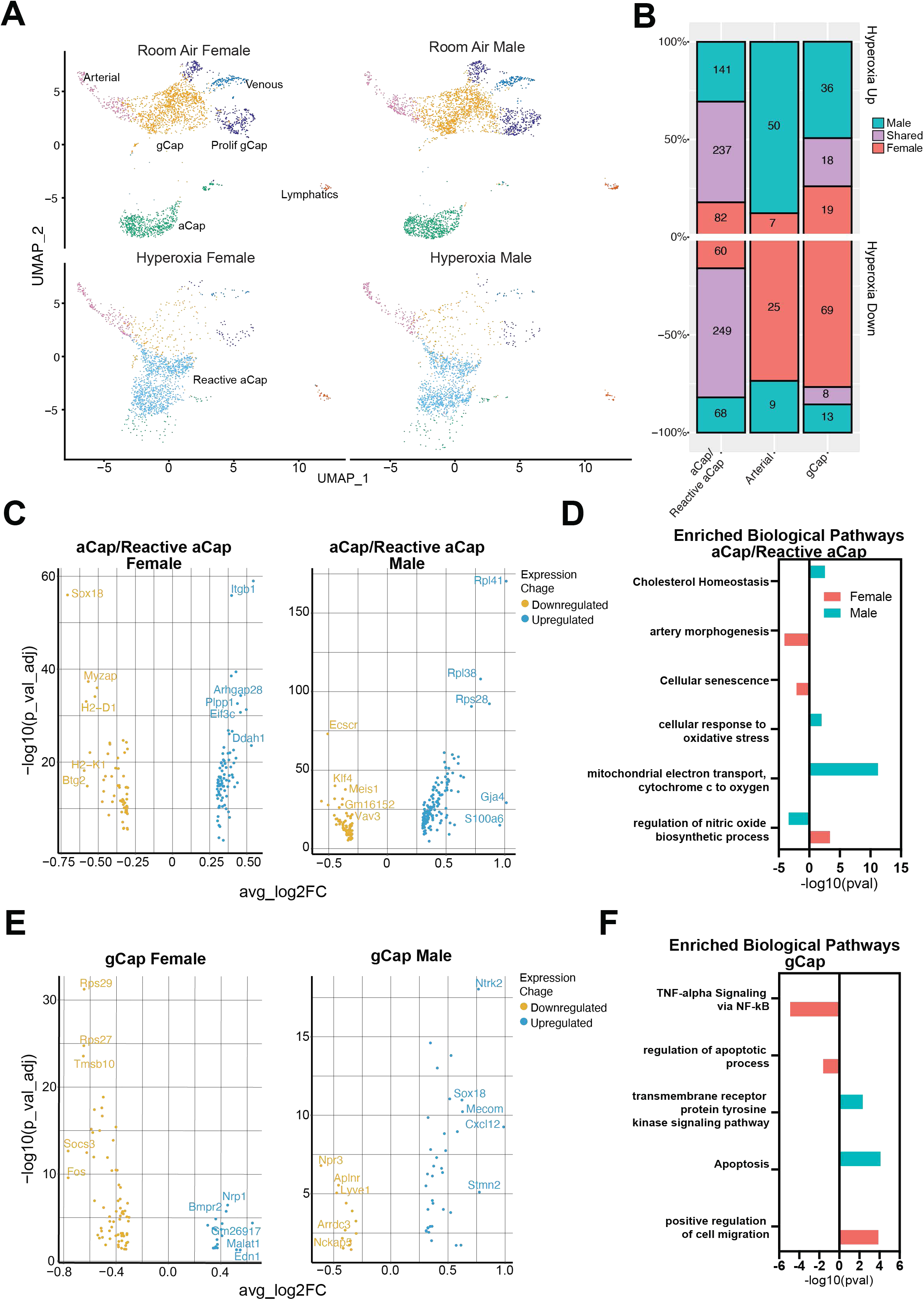
Sex Specific Response to Hyperoxia in Endothelial Cells. **A)** UMAP of sequenced lung endothelial cells identified seven distinct clusters split by sex and experimental condition (room air and hyperoxia). **B)** Number of differentially expressed genes (upregulated on top and downregulated at the bottom) in response to hyperoxia that are either shared between male and female (purple), unique in male (blue) or unique in female (pink) aCaP, arterial and gCaP lung endothelial cells **C)** Volcano plots showing up- and down- regulated in the female and male aCaP/reactive aCaP endothelial cells upon exposure to hyperoxia. **D)** Sex-specific enriched biological pathways (blue: male, pink: female) in aCaP/reactive aCaP lung endothelial cells. **E)** Volcano plots showing up- and down- regulated in the female and male gCaP endothelial cells upon exposure to hyperoxia. **F)** Sex-specific enriched biological pathways (blue: male, pink: female) in gCaP lung endothelial cells.

Among gCaP lung capillary endothelial cells, *CxCl12* was upregulated in male, while neuropilin-1 (*Nrp1)* and bone morphogenetic receptor 2 (*Bmpr2)* were upregulated in females (**Figure 3E**). *CxCl12* plays an important role in angiogenesis through interaction with its receptors CXCR4 or CXCR7 (53–55). Autocrine signaling in lung endothelial cells leads to the development of pulmonary hypertension (56, 57). *Nrp1* mediated VEGF signaling in endothelial cells and play a key role in angiogenesis. *Nrp1* also attenuates senescence in endothelial cells(58). Among sexually dimorphic pathways in gCaPs, TNF-signaling was downregulated, while cell migration was upregulated in females. Transmembrane tyrosine receptor kinase was upregulated in males. Apoptosis was modulated in opposite directions in males and females (**Figure 3F**). Differentially expressed genes in male and female endothelial cell sub-populations are listed in **Supplemental Table 2.**

### Lung Epithelial gene expression changes in response to hyperoxia

Lung epithelial cells comprised of AT1, AT2, Lyz1+ AT2 and ciliated epithelial cells (**Figure 4A**) and their relative contribution to the lung epithelial cells in room air and hyperoxia is shown in **Figure 4B and 4C**. The number of cells analyzed and the number of upregulated and downregulated genes in AT1, AT2 and LyZ1+ AT2 cells is shown in **Figure 4D**. p21 (*Cdkn1a*) was upregulated in hyperoxia exposed AT1 and AT2 cells. Peroxiredxin (*Prdx*) 6 was upregulated in AT1 cells, which is induced in the alveolar epithelium under oxidative stress and has an antioxidant role (59). *Igfbp2* was downregulated in the AT1 cells, which is a marker for postnatal AT1 cells and increases following pneumonectomy during lung regeneration. *Igfbp2* positive AT1 cells are terminally differentiated (60). Biological pathways related to oxidative phosphorylation, apoptosis and neutrophil mediated immunity were upregulated, while TNF-signaling via NF-kappa B, hypoxia, and the MAP kinase signaling pathway were downregulated (**Figure 4E**).

**Figure 4.**
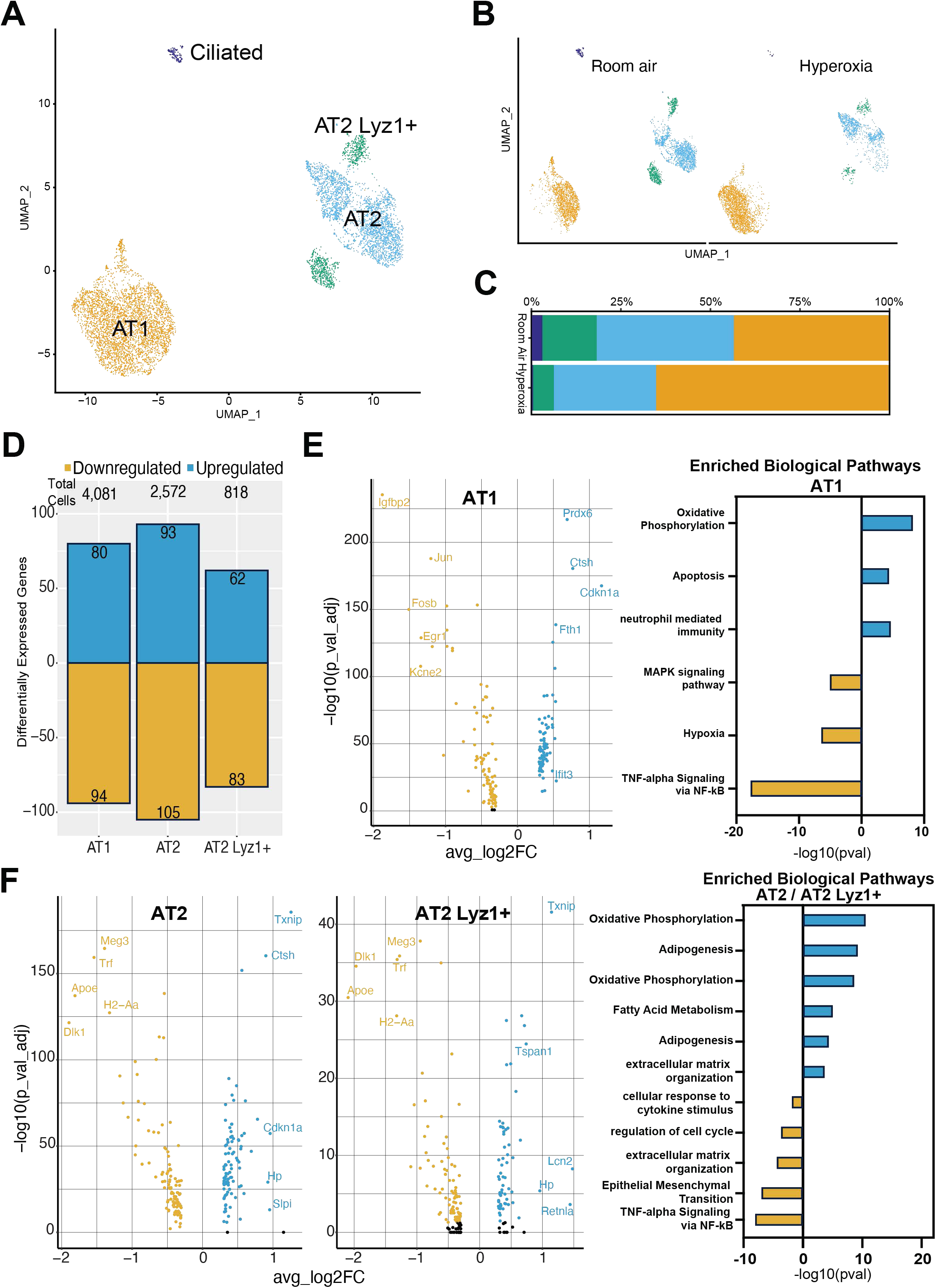
Gene Expression Changes in Epithelial Cells in Response to Hyperoxia. **A)** UMAP of sequenced lung epithelial cells identified four distinct clusters. **B)** UMAP of lung endothelial cells showing the four distinct clusters in room air and hyperoxia. **C)** Changes in the relative contribution of different lung epithelial cell sub-populations in room air and hyperoxia. **D)** Number of cells sequenced and the number of up- and down-regulated genes in AT1, AT2 and AT2 Lyz 1+ lung epithelial cells upon exposure of the neonatal lung to hyperoxia. **E)** Volcano plots showing the differentially expressed genes in AT1 cells in response to hyperoxia and enriched biological pathways. F) Volcano plots showing the differentially expressed genes in AT2 and AT2 Lyz1+ cells in response to hyperoxia and enriched biological pathways.

Thioredoxin interacting protein (*Txnip*) was induced in the AT 2 and AT2 Lyz1+ cells upon exposure to hyperoxia. Overexpression of *Txnip* decreased HIF activity and VEGF expression in the murine lung independent of its ability to bind thioredoxin (61). *Apoe* was downregulated in the hyperoxia exposed AT2 and AT2 Lyz1+ cells which is protective in lung disease models through its anti-inflammatory and anti-oxidant effects (62). Oxidative phosphorylation and adipogenesis were upregulated, while extracellular matrix organization and epithelial to mesenchymal transition were among the downregulated pathways in AT2 and AT2 Lyz1+ cells (**Figure 4F**). Differentially expressed genes in all epithelial cells are listed in **Supplemental Table 3.**

### Sex specific response to hyperoxia in epithelial cells

The lung epithelial cells in room air and hyperoxia in the male and female PND7 murine lung are shown in **Figure 5A**. The number of up- and down-regulated differentially expressed genes show remarkable sex-specific modulation of the cellular transcriptome with most genes showing a sex-specific expression with little overlap (**Figure 5B**). ATPase Na+/K+ Transporting Subunit Beta 1(*Atp1b1*) was upregulated in the female AT1 cells (**Figure 5C**), which is crucial for alveolar fluid clearance (63), and expression is decreased in lung fibrosis (64). *Egr1 (early growth response gene 1)* and *Fosb* are downregulated in the male AT1 cells upon exposure to hyperoxia, both of which are immediate early genes and inhibit cell proliferation (65, 66). Surfactant protein D (*Sftpd*) was upregulated in the female AT2 cells (**Figure 5C**) which is a hydrophilic collectin molecule secreted by Type 2 alveolar epithelial cells and plays an anti-inflammatory role by regulating lung macrophage activity (67). Secretory leukocyte proteinase inhibitor (*Slpi*) was increased in male AT2 cells, which is an anti-protease and acts on neutrophil elastase (68, 69). Cytochrome c oxidase subunit 6c (*Cox 6c*) was upregulated in male AT2 Lyz1+ cells (**Figure 5C**), which plays a crucial role in oxidative phosphorylation and may play a role in the metabolic changes in these cells in response to hyperoxia (70). Sex-specific biological pathways enriched from genes that were exclusively differentially regulated in male and females in AT1 and AT2 cells are highlighted in **Figure 5D**. Angiogenesis was downregulated in male, while apoptosis, inflammatory response and interferon-gamma response were downregulated in female AT1 cells. Aerobic electron transport chain was upregulated in male, while ECM-receptor interaction upregulated in female AT1 cells. Aerobic electron transport chain was upregulated, while xenobiotic metabolism was downregulated in male AT2 cells, while coagulation, myogenesis and cytokine mediated signaling pathway was downregulated in female AT2 cells. TNF-alpha signaling via NF-kappa B and p53 pathway were downregulated in female AT2 Lyz1+ cells (**Figure 5D**). Differentially expressed genes in male and female lung epithelial cell sub- populations are listed in **Supplemental Table 3.**

**Figure 5.**
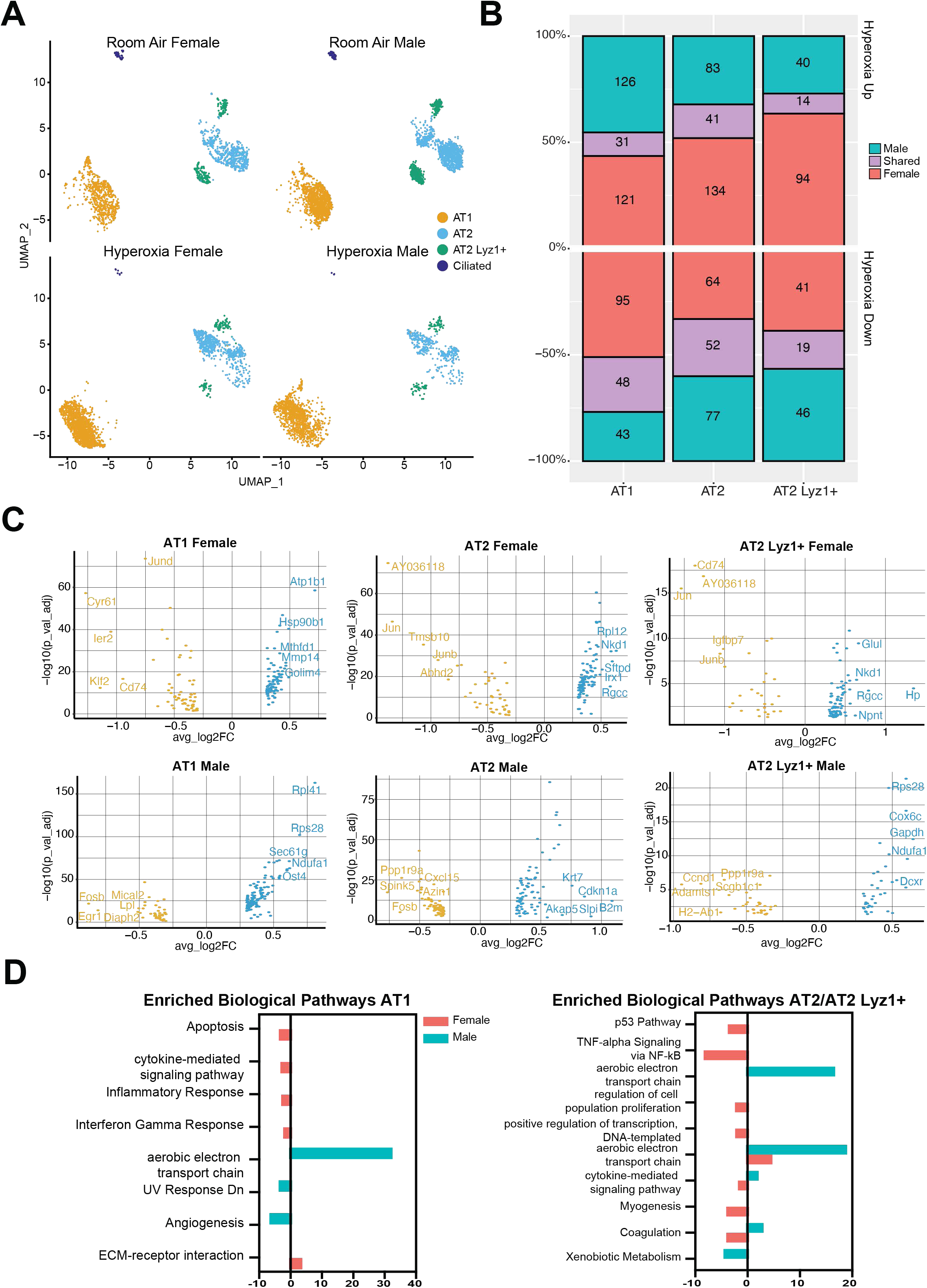
Sex Specific Response to Hyperoxia in Epithelial Cells. **A)** UMAP of sequenced lung epithelial cells identified four distinct clusters split by sex and experimental condition (room air and hyperoxia). **B)** Number of differentially expressed genes (upregulated on top and downregulated at the bottom) in response to hyperoxia that are either shared between male and female (purple), unique in male (blue) or unique in female (pink) AT1, AT2 and AT2 Lyz1+ lung epithelial cells **C)** Volcano plots showing up- and down- regulated in the lung AT1, AT2 and AT2 *Lyz1+* lung epithelial cells in female and male lungs upon exposure to hyperoxia. **D)** Sex-specific enriched biological pathways (blue: male, pink: female) in AT1 and AT2 lung epithelial cells.

### Neonatal hyperoxia exposure leads to significant changes in cell state in lung immune cells

Sub clustering of 13,103 immune cells identified 12 distinct populations (**Figure 6A**). UMAP plots of the immune cell sub-populations in room air and hyperoxia are shown in **Figure 6B** and % contribution to the total immune cells under room air and hyperoxia is shown in **Figure 6C.**

**Figure 6.**
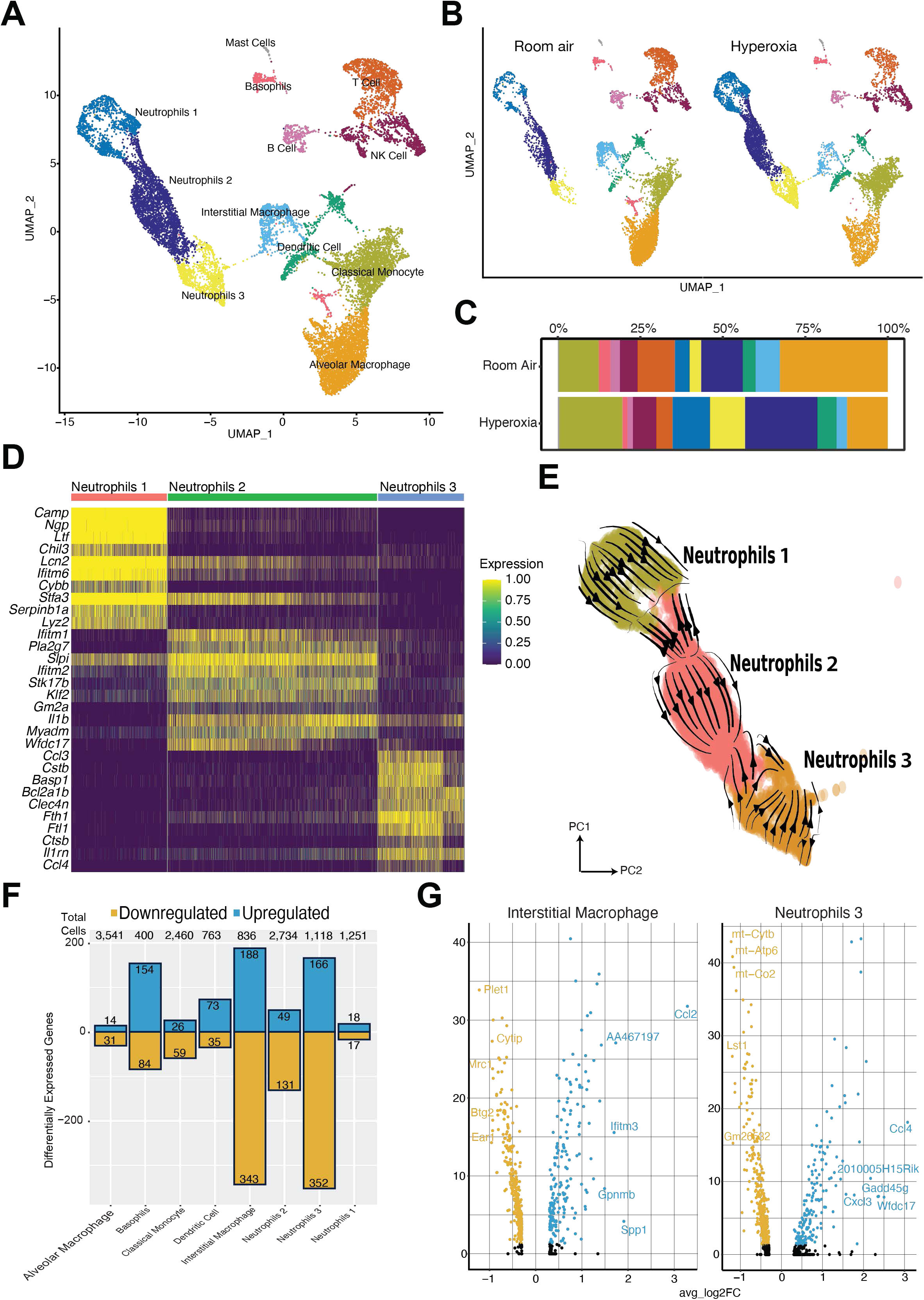
Gene Expression Changes in Immune Cells in Response to Hyperoxia. **A)** UMAP of sequenced lung immune cells identified twelve distinct clusters. **B)** UMAP of lung endothelial cells showing the twelve distinct clusters in room air and hyperoxia. **C)** Changes in the relative contribution of different lung immune cell sub-populations in room air and hyperoxia. **D)** Gene expression similarities and differences between neutrophils 1, 2 and 3. **E)** Trajectory analysis of lung neutrophils showing neutrophils 1 and 3 arise from neutrophil 2. Neutrophils 1 and 3 show gene signatures seen in PMN-MDSCs and activated PMN-MDSCs respectively. **F)** Number of cells sequenced and the number of up- and down-regulated genes in lung immune cells upon exposure of the neonatal lung to hyperoxia. **G)** Volcano plots showing the differentially expressed genes in interstitial macrophages and neutrophils 3 in the neonatal lung upon exposure to hyperoxia.

We identified a population of neutrophils enriched for markers associated with PMN- myeloid derived suppressor cells (PMN-MDSCs) (neutrophils 1 and 3) (26). PMN-MDSCs possess immunosuppressive properties and suppress the function of T-, B- and NK cells and are thought to be pathologically activated. They are considered to be pro-tumorigenic due to their immune-suppressive functions and contribute to pathologic tissue remodeling (71). Neutrophils 3 in our study expand under hyperoxic conditions. This cell cluster is similar to the neutrophil cluster reported by Hurskainen *et al* (*24*) after the neonatal murine lung was exposed to hyperoxia with higher expression of *Basp1* and *CCl4.* Differences in gene expression between these three neutrophil populations are shown in **Figure 6D.** Upon exposure to hyperoxia, neutrophils 3 enriched for *Ccl4, Ccl3, CxCl4, Basp1, Ninj1, Hilpda, Gadd45b, Atf3 and Mif* associated with activated PMN- myeloid derived suppressor cells (26). Neutrophils 1 showed enrichment for genes associated with immature neutrophils and PMN-MDSCs (*Ngp, Ltf, Cybb*)(26). Trajectory analysis using RNA velocity showed that neutrophils 1 and 3 arise from neutrophils 2 (**Figure 6E**). Supplemental table showing enriched genes in these three neutrophil populations are included in **Supplemental Table 4.**

The number of cells sequenced and up- and down-regulated genes in each immune cell population is shown in **Figure 6F.** The interstitial macrophage and the Type 3 neutrophils showed the greatest number of differentially expressed genes among the immune cells following exposure to hyperoxia. Interstitial macrophages showed high expression of *Spp1* **(Figure 6G)** or osteopontin (72). Interstitial macrophages are sources of *Spp1*, which functions as a pro-fibrotic mediator in lung diseases (73, 74). Type 3 neutrophils enrich for the chemokines *CCl4* and *CxCl3.* These are characteristic of activated PMN-MDSCs(26). Leukocyte specific transcript-1 (*Lst1*) was one of downregulated genes in Type 3 neutrophils **(Figure 6G)**. Interestingly, this finding was also reported among lung neutrophils in an IL-beta mediated pneumonitis model (75).

The common biological pathways up- and down-regulated in male and female lungs in interstitial macrophages (**Supp Figure 2A)** and Type 3 neutrophils (**Supp Figure 2B)**. Biological pathways enriched in interstitial macrophages included interferon gamma response, complement, oxidative phosphorylation, necroptosis and cellular response to interkeukin-1. Neutrophils 3 showed upregulation of TNF-alpha signaling via NF-kappaB, inflammatory response, IL-2/STAT 5 signaling, apoptosis and neutrophil degranulation, while adipogenesis was downregulated. Differentially expressed genes in all immune cells are listed in **Supplemental Table 5.**

### Sex-specific changes in the lung immune cells in response to neonatal hyperoxia

The lung immune cells show remarkable sex-specific differences in their transcriptional state upon exposure to hyperoxia. The UMAP plots of the lung immune cells in male and female lungs in room air and hyperoxia are shown in **Figure 7A**. The number of unique and shared differentially expressed genes in the immune cell subpopulations are shown in **Figure 7B.** Basophils, B cells, neutrophils 3, and classical monocytes are predominated by the female response, while interstitial macrophages show a predominant response in males. Volcano plots showing the DEGs in male and female alveolar macrophages, interstitial macrophages and neutrophils 3 are shown in **Figure 7C**. DEGs in female basophils and B cells are shown in **Figure 7D**. The top 2 upregulated genes in basophils in the female lung (**Figure 7D**) upon exposure to hyperoxia were *Mcpt8 (mast cell protease 8)* (*76*) and *Prss34 (protease serine 34)*(*77*) both of which are specific to basophils. Upregulated pathways in basophils included interferon gamma response and Myc targets, while TNF-alpha, IL-6, PI3- Akt signaling and cellular response to TGF-beta were among the downregulated pathways **(Figure 7E).**

**Figure 7.**
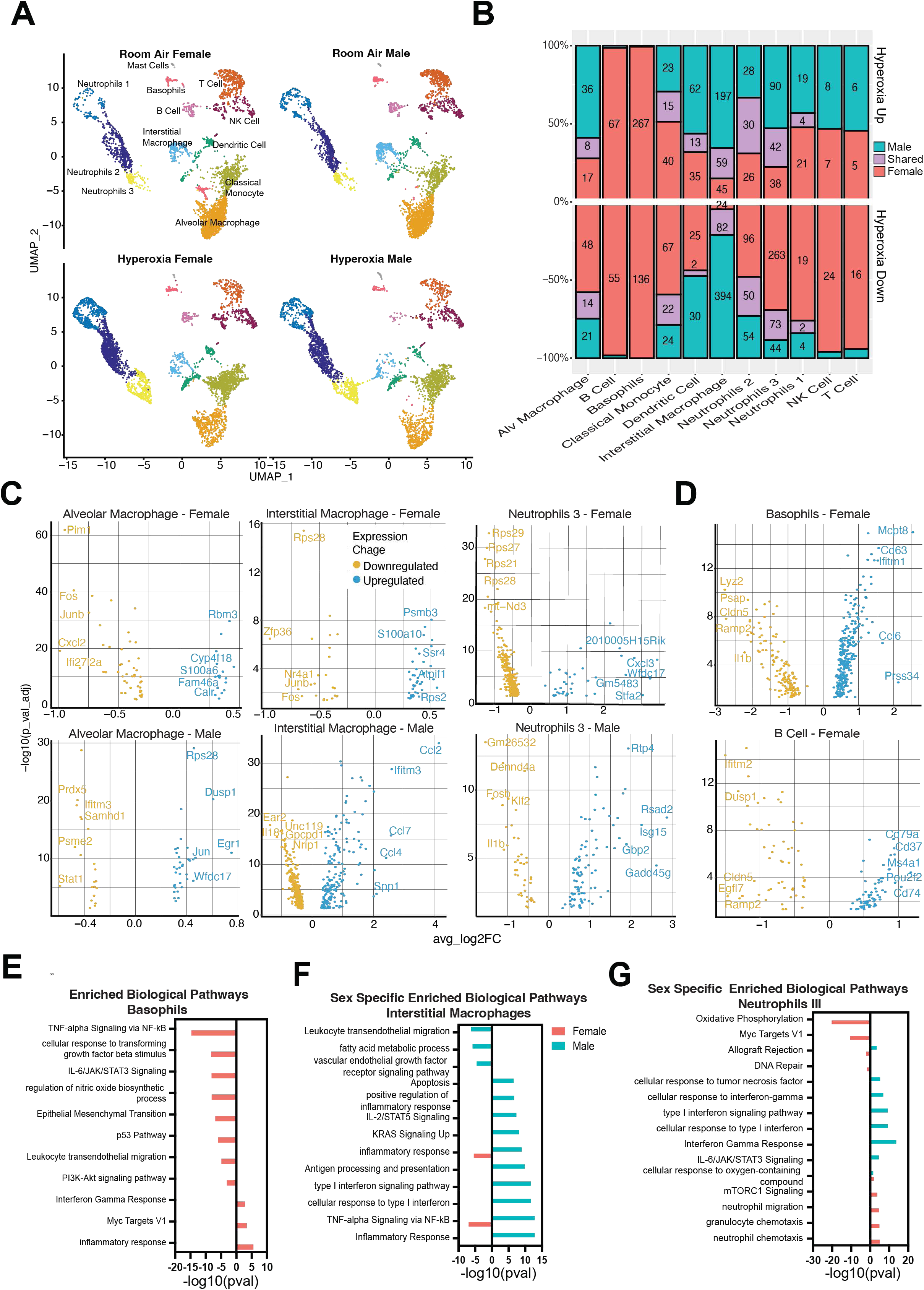
**Sex Specific Response to Hyperoxia in Immune Cells**. **A)** UMAP of sequenced lung immune cells identified twelve distinct clusters split by sex and experimental condition (room air and hyperoxia). **B)** Number of differentially expressed genes (upregulated on top and downregulated at the bottom) in response to hyperoxia that are either shared between male and female (purple), unique in male (blue) or unique in female (pink) lung immune cells **C)** Volcano plots showing up- and down- regulated genes in female and male alveolar macrophages, neutrophils 3 and interstitial macrophages upon exposure to hyperoxia. **D)** Volcano plots showing up- and down- regulated genes in female basophils and B cells. Sex- specific enriched biological pathways (blue: male, pink: female) in **E)** basophils, **F)** Interstitial macrophages and **G)** neutrophils 3.

*CCl4* or macrophage inflammatory protein-1 beta was upregulated in the male IMs, that plays a role in leukocyte recruitment following injury (**Figure 7C**) (78). Biological pathways upregulated in male lung interstitial macrophages upon exposure to hyperoxia included type 1 interferon signaling, inflammatory response, antigen processing and presentation, IL- 2/STAT 5 signaling and TNF-alpha signaling via NF-kappa B. On the contrary inflammatory response was downregulated in female interstitial macrophages (**Figure 7F**). *Gadd45g* was upregulated in male neutrophils 3 that plays a role DNA repair (**Figure 7C**) (79, 80). Female specific pathways downregulated in neutrophils 3 included oxidative phosphorylation, DNA repair, Myc targets, while neutrophil chemotaxis and mTORC1 signaling were upregulated. Interferon-gamma response, IL-6 signaling, cellular response to type 1 interferon and TNF were upregulated in male neutrophils 3 (**Figure 7G**). *S100a4* was upregulated in male classical monocytes, that leads to increased production of inflammatory mediators from monocytes (81), while *CxCl2* was downregulated in females in classical monocytes, which recruits neutrophils and leads to neutrophil accumulation (82) (**Supp Figure 2C**). Classical monocytes in male lungs showed upregulation of neutrophil mediated immunity, PI3-Akt signaling and complement, while in females MAPK signaling pathway, inflammatory response, TNF-alpha signaling via NF-kappaB and regulation of apoptosis were downregulated (**Supp Figure 2D**). Differentially expressed genes in male and female lung immune cell sub-populations are listed in **Supplemental Table 5.**

### Cell-cell communication

To infer, visualize and analyze intercellular communications from the scRNA-seq data we used the Cell Chat tool (83). We interrogated whether the cell-cell interaction between the cell types was significantly changed and if the major sources and targets were changed between room air and hyperoxic conditions. We analyzed the cell-cell communication in the endothelial, epithelial and immune cell subpopulations and also studied the interactions between epithelial-endothelial and endothelial-immune subpopulations.

*Novel ligand-receptor pairs and the role of the distal lung endothelium in neonatal hyperoxia* The cell-cell communication between the different endothelial cell subpopulations is enhanced, but the interaction strength is decreased compared to normoxia **(Supp Figure 3A).** Under room air conditions, among all endothelial cell subpopulations, the aCaPs have significant outgoing and incoming signaling to and from gCaPs and proliferating gCaPs (as demonstrated by the edge width of the ligand-receptor pairs) and serve as the dominant communication hub (**Figure 8A**). Strikingly, upon exposure to hyperoxia, the differential number of incoming and outgoing signaling networks are increased from the reactive aCaP population while the signaling networks among other endothelial cell subpopulations is decreased compared to normoxia (**Figure 8 A, B**). There was a strong component of autocrine signaling in the reactive aCaPs (**Figure 8B**). Within the endothelial cell sub- population, we identified the global communication patterns that connect cell groups with signaling pathways in the context of outgoing or incoming signaling under room air and hyperoxia conditions (**Figure 8C-D**). We uncovered three outgoing and incoming signaling patterns under hyperoxia. There are notable differences and similarities between the communication patterns in room air and hyperoxia. The communication patterns in room air are shown in **Figure 8C**, where the outgoing signaling shows four patterns with the aCaPs, gCaPs, arterial/venous and lymphatic endothelial showing distinct outgoing signaling pathways. Under hyperoxia, a large portion of the outgoing signaling from the distal lung endothelium (with gCaPs, aCaPs and reactive aCaPs) is characterized by pattern #1, which represents multiple pathways, including but not limited to *Kit*, *Apelin*, *Semaphorin* and *collagen* pathways. The lymphatic and arterial/venous endothelial cells have distinct outgoing signaling pathways. The incoming signaling patterns in hyperoxia grouped the gCaP/proliferating gCaP (pattern #3), arterial/venous/lymphatic (pattern #1) and the aCaPs (pattern #2) in three groups (**Figure 8D**). *Collagen*, *Kit*, *Semaphorin* (*Sema) 7* and protein tyrosine phosphatase receptor type M (*Ptprm)* were the autocrine-acting pathways in the gCaP/proliferating gCaP population. Since the reactive aCaPs population was significantly increased in the hyproxia-exposed PND7 lung, we focused on the incoming and outgoing signaling pathways in this cell subpopulation. *Tweak* (*Tnf superfamily member 12*) and *Ptprm* pathways were identified in the incoming and outgoing signaling pathways in this population of cells respectively **(Figure 8E-F).** *Tweak* belongs to the TNF ligand family and is a ligand for the *Fn14/TweakR* receptor. It plays a role in angiogenesis by modulating endothelial cell proliferation and migration and has been shown to promote oxidative stress in endothelial cells (84–87). *Ptprm* (protein tyrosine phosphatase receptor type M) is highly expressed in the lung and expressed in the lung endothelium. It binds directly to VE-cadherin, modulates its phosphorylation, and maintains barrier integrity (88, 89) . The Ptprm pathway was not among the inferred networks under room air conditions in lung endothelial cells. Network centrality analysis of the inferred *Ptprm* signaling network identified that reactive aCaPs are the most prominent sources and dominant mediators for this ligand and has a predominant autocrine and paracrine effect on gCaPs and proliferating gCaPs, with no discerned effects on other endothelial cells. The TGF-beta network between room air and hyperoxia, displayed some striking differences. The aCaPs are the predominant targets under room air and hyperoxia (**Figure 8G-H**). TGF-beta 1 was the main ligand with expression from all endothelial cells under room air and hyperoxia. However, TGF-beta 2 was induced in reactive aCaPs and lymphatic endothelial cells upon exposure to hyperoxia (**Figure 8I**). These two isoforms have different tissue specific expression(90, 91) and may have distinct biological functions in disease pathophysiology (92).

**Figure 8.**
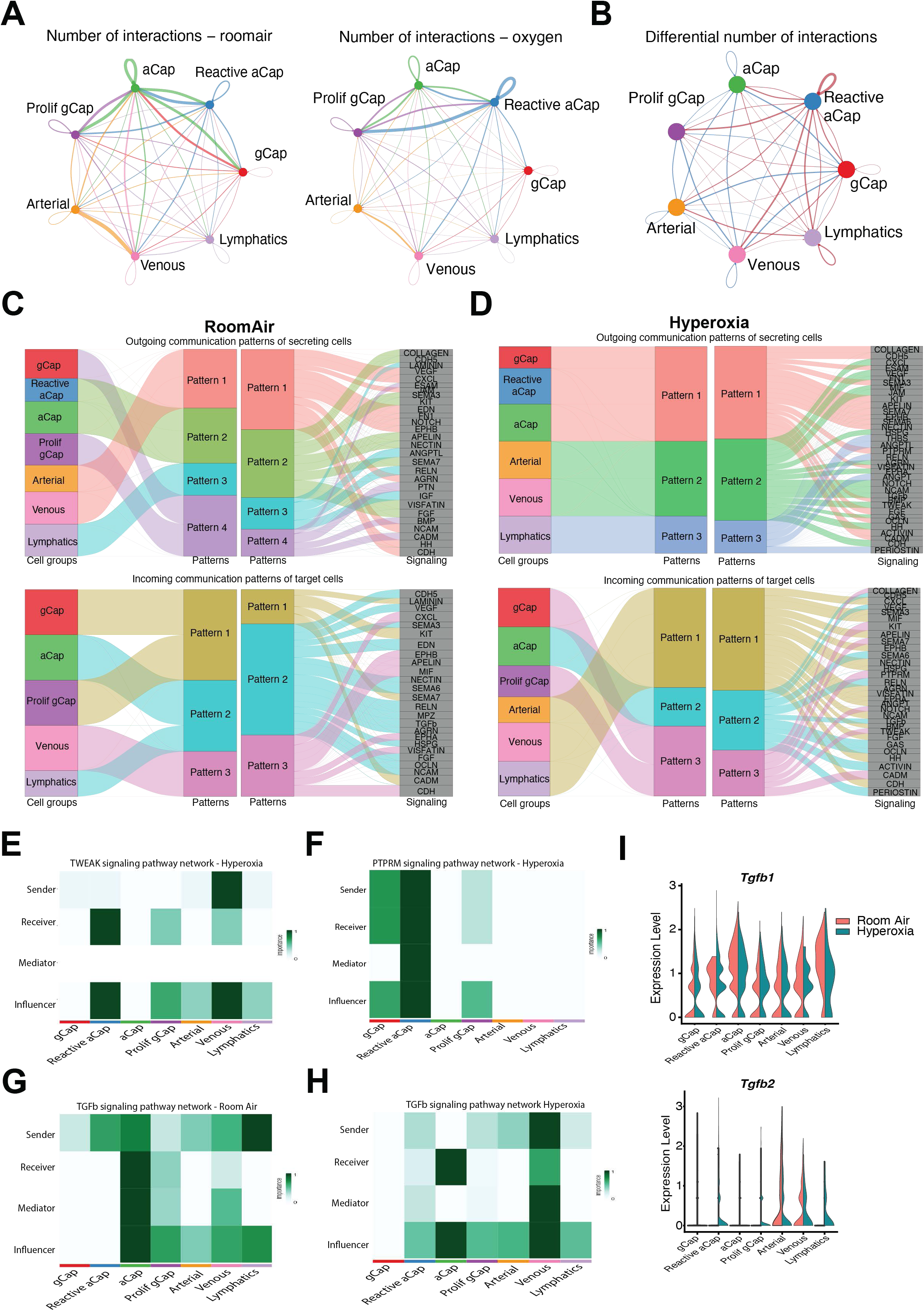
Intercellular Communication Changes in Response to Hyperoxia in Lung Endothelial Cells. **A)** Circle plot showing the number of interactions between different endothelial cell subpopulations in room air and hyperoxia. Edge colors are consistent with the sources as sender, and edge weights are proportional to the interaction strength. Thicker edge line indicates a stronger signal. Circle sizes are proportional to the number of cells in each cell group. **B)** Circle plot showing the number of differential interactions between different endothelial cell sub-populations in room air (blue) and hyperoxia (red). The inferred outgoing patterns of secreting cells and incoming communication patterns of target cells among the lung endothelium, in room air **C)** and hyperoxia **D)**. The thickness of the flow indicates the contribution of the cell group or signaling pathway to each latent pattern. Heatmap showing the relative importance of each cell group based on the network centrality measures of **E)** *Tweak* signaling network in hyperoxia, **F)** *Ptprm* signaling network in hyperoxia, *TGF-beta* signaling in room air **(G)** and hyperoxia **(H)**. Violin plots showing expression of *Tgfb1* and *Tgfb2* **(I)** in lung endothelial cells in room air and hyperoxia.

### Neonatal hyperoxia alters cell to cell communication among lung epithelial cells

The number of cell-cell communication networks is decreased, but their strength increased in hyperoxia compared to normoxia among lung epithelial cells **(Supp Fig 3B)**. Under both room air and hyperoxic conditions, AT1 cells enrich in outgoing signaling pathways while AT2 cells enrich in incoming signaling pathways **(Supp Fig 3C).** However, the number of cell-cell communication from the AT1 cells was decreased in hyperoxia compared to room air conditions **(Figure 9 A-B).** The communication patterns between alveolar epithelial cells in room air and hyperoxic conditions are highlighted in **Figures 9 (C-D)**. Under hyperoxia, AT1 cells are the senders for most of the outgoing signaling networks (pattern #1).

**Figure 9:**
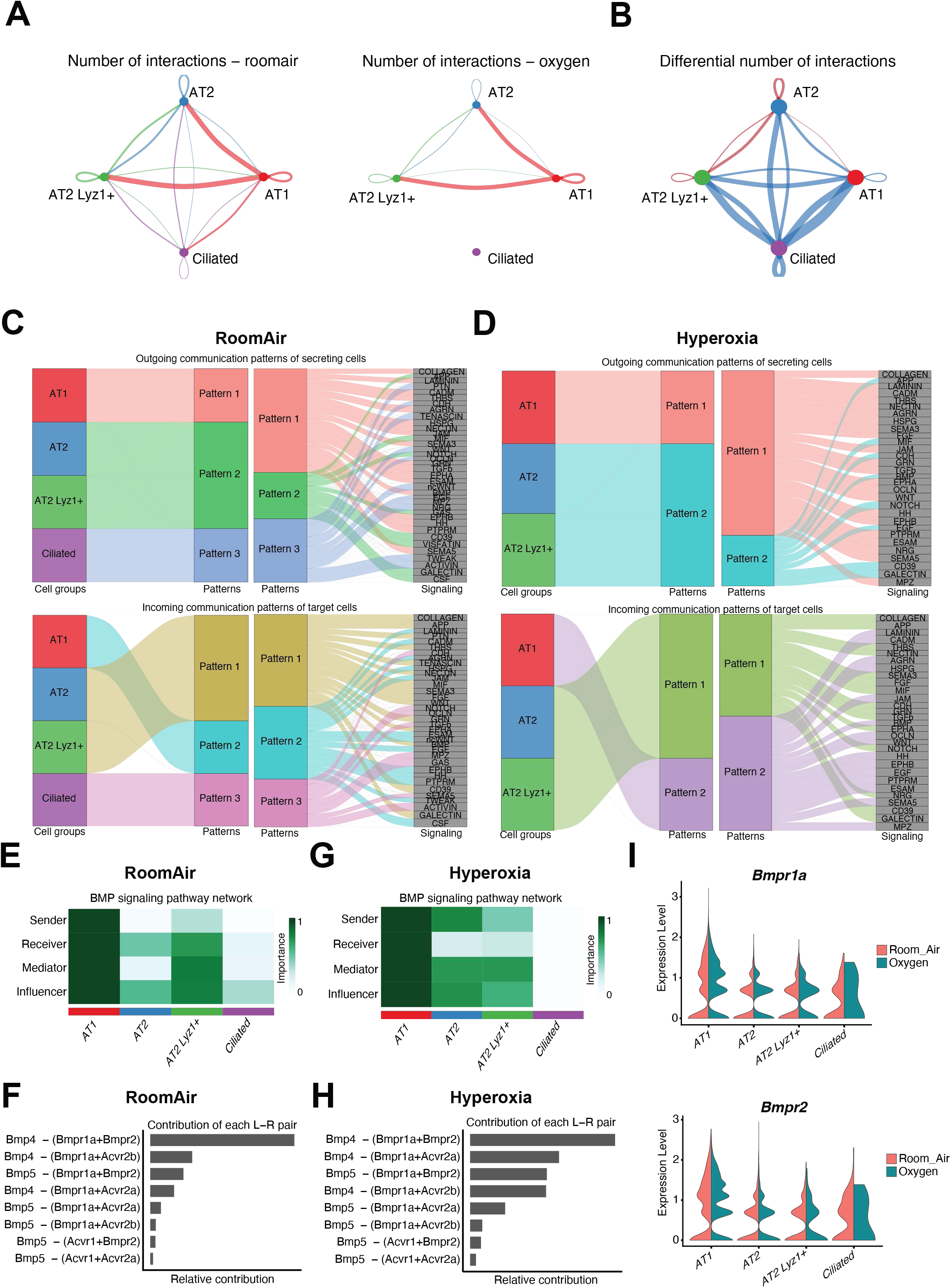
Intercellular Communication Changes in Response to Hyperoxia in Lung Epithelial Cells. **A)** Circle plot showing the number of interactions between different lung epithelial cell subpopulations in room air and hyperoxia. Edge colors are consistent with the sources as sender, and edge weights are proportional to the interaction strength. Thicker edge line indicates a stronger signal. Circle sizes are proportional to the number of cells in each cell group. **B)** Circle plot showing the number of differential interactions between different endothelial cell sub-populations in room air (blue) and hyperoxia (red). The inferred outgoing patterns of secreting cells and incoming communication patterns of target cells among the lung epithelial cells, in **C)** room air and **D)** hyperoxia. The thickness of the flow indicates the contribution of the cell group or signaling pathway to each latent pattern. Heatmap showing the relative importance of each cell group based on the network centrality measures of **E)** *Bmp* signaling network in room air and **F)** relative contribution of each ligand-receptor pair in room air to the overall communication network of *Bmp* signaling pathway. Heatmap showing the relative importance of each cell group based on the computed four network centrality measures of **G)** *Bmp* signaling network in hyperoxia and **H)** relative contribution of each ligand-receptor pair in hyperoxia to the overall communication network of *Bmp* signaling pathway **I)** Violin plots showing expression of *Bmpr1a* and *Bmpr2* in lung epithelial cells in room air and hyperoxia.

The bone morphogenetic protein (BMP) signaling network, under room air conditions originates from the AT1 cells (sender) with autocrine and paracrine effects on the AT1 and AT2 cells (receiver). Under hyperoxia, the AT1 cells are the main receivers with AT1 and AT2 cells becoming the ligand sources (**Figure 9 E-F**). *Bmp4* was the main ligand, with *Bmpr2* and *Bmpr1a* were the inferred receptors **(Figure 9 G-H).** *Bmpr1a* and *Bmpr2*expression was increased in AT1 cells after exposure to hyperoxia **(Figure 9I).** The role of BMP signaling in the alveolar niche has been highlighted in a pneumonectomy model with increased BMP signaling enhancing the differentiation of AT2 cells to AT1 cells (93). In primary cultures of mouse AT-2 cells, however, recombinant BMP4 inhibited trans differentiation of AT2 to AT1 cells (94).

#### Cell-cell communication patterns among lung immune cells are altered by neonatal hyperoxia

The number of cell-cell communication networks among the immune cells are increased in hyperoxia compared to normoxia, while the interaction strength decreased **(Supp Figure 4A).** Classical monocytes and interstitial macrophage enrich in pathways related to outgoing signaling both under room air and hyperoxia while dendritic cells enrich in incoming signaling networks. Basophils emerge as a cell subpopulation with increase in outgoing interaction strength upon exposure to hyperoxia **(Supp Figure 4B).** Interstitial macrophage and dendritic cells also show strong autocrine signaling in the hyperoxia exposed lung **(Figure 10A).** The communication patterns between immune cells in the lung in room air and hyperoxia are shown in **Figure 10B-C.** Interstitial macrophages have a distinct set of outgoing and incoming signaling patterns both in room air and hyperoxia. Interestingly, enriched among the outgoing signaling patterns in basophils in hyperoxia is GM colony stimulating factor (Csf), with macrophages and dendritic cells as receivers (**Figure 10D**). *Csf1* expression is increased in basophils, while *Csf1r* expression is increased in lung dendritic cells upon exposure to hyperoxia (**Figure 10D**). Basophils are imparted a unique transcriptional signature in the lung microenvironment partly by GMCSF and polarize the alveolar macrophages towards an anti-inflammatory state (95). The fibronectin signaling pathway was activated in hyperoxia with classical monocytes and interstitial macrophages as sources and CD44 as the receptor **(Figure 10E).** Increased expression of fibronectin in lung macrophages has been shown in lung diseases and upon exposure to hyperoxia (96, 97).

**Figure 10:**
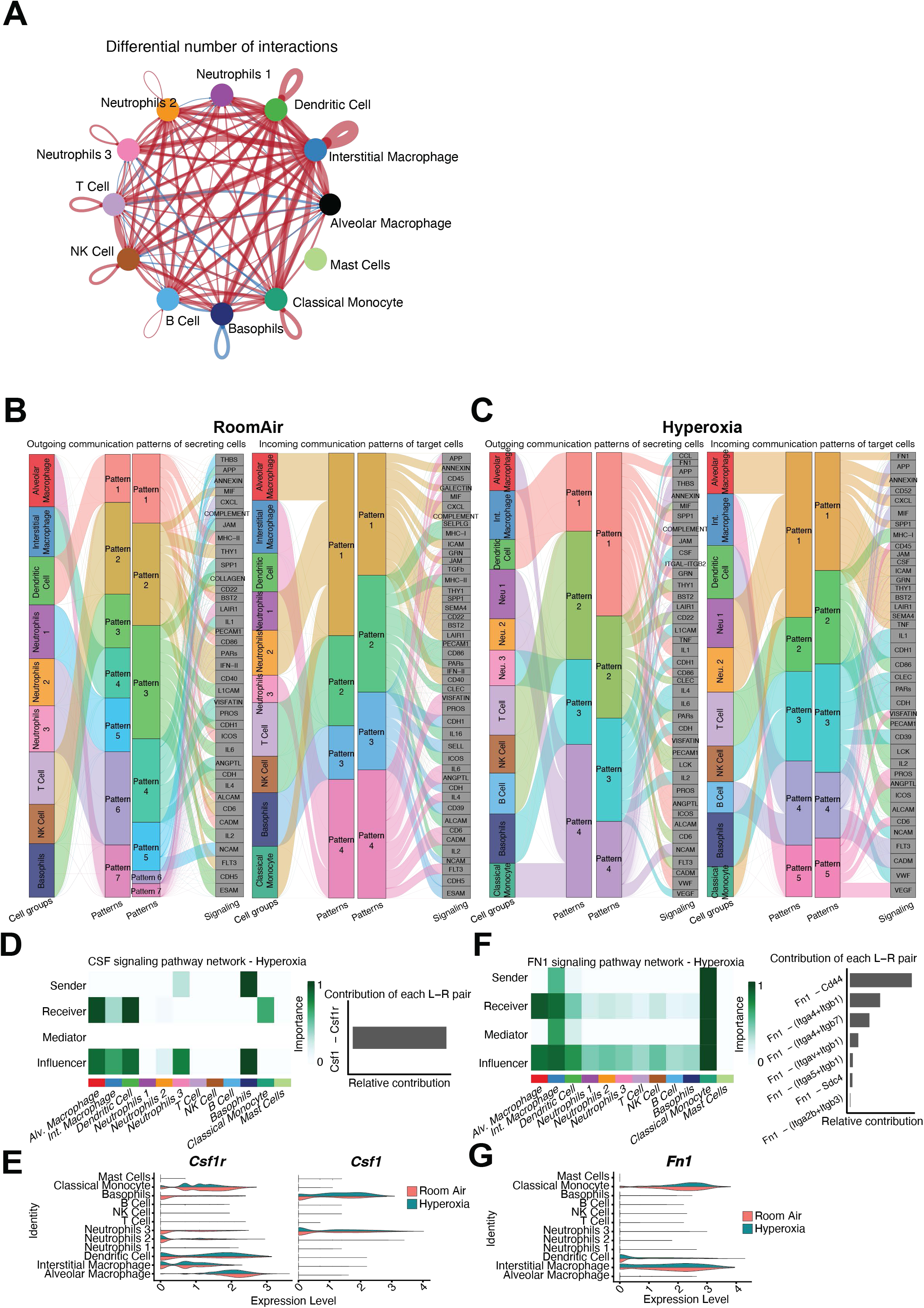
Intercellular Communication Changes in Response to Hyperoxia in Lung Immune Cells. **A)** Circle plot showing the number of differential number of interactions between different lung immune cells in room air (blue) and hyperoxia (red). The inferred outgoing patterns of secreting cells and incoming communication patterns of target cells among the lung immune cells, in room air **B)** and hyperoxia **C)**. The thickness of the flow indicates the contribution of the cell group or signaling pathway to each latent pattern. Heatmap showing the relative importance of each cell group based on the network centrality measures of **D)** *Csf1* signaling network in hyperoxia, and relative contribution of each ligand- receptor pair in room air to the overall communication network of *Csf1* signaling pathway, and **E)** violin plots showing expression of *Csf1* and *Csf1r* in lung immune cells in room air and hyperoxia. Heatmap showing the relative importance of each cell group based on the computed four network centrality measures of **F)** *Fn1* signaling network in hyperoxia, and relative contribution of each ligand-receptor pair in room air to the overall communication network of *Fn1* signaling pathway, and **G)** violin plots showing expression of *Fn1* in lung immune cells in room air and hyperoxia.

### Neonatal hyperoxia decreases and alters ligand-receptor interactions between endothelial and epithelial cells

The number of inferred interactions and the interaction strength decreases upon exposure to hyperoxia **(Supp Figure 4C)**. aCaPs, reactive aCaPs and AT1 cells showed the highest outgoing interaction strength, while the AT2 cells the highest incoming interaction under normoxia and hyperoxia **(Supp Figure 4D).** Reactive aCaP cells display the most differential number and strength of both outgoing and incoming signaling networks from AT1, AT2 and AT2Lyz1+ cells **(Figure 11 A-B).** The communication patterns between epithelial and endothelial cells in room air and hyperoxia are shown in **Figure 11 C-D.** Under room air, most of the outgoing signaling was dominated by pattern #2 (from AT1 epithelial cells) and pattern#3 was from ciliated epithelial cells. Pattern #2, in room air from AT1 epithelial cells included VEGF with aCaP cells as receivers. Under hyperoxia, four predominant outgoing patterns were observed with pattern #1 (reactive aCaPs/ aCaPs) and pattern #2 (AT1 cells) being the major secreting cells. The *activin* signaling network (ligand receptor pair: Inhba- (Acvr1b+Acvr2a)) signals from the reactive aCaPs to AT2 and AT2 Lyz1+ cells (**Figure 11 E**). Endothelial activin-A signaling accelerated the development of pulmonary hypertension (98). Activin signaling also exacerbated pathology in murine models of idiopathic pulmonary fibrosis (99–101), and overexpression led to inflammation and alveolar cell death (102). *Tweak* (*Tnf superfamily member 12*) is another inferred endothelial to epithelial communication with the ligand from the gCaPs, aCaP, and arterial endothelial cells and signaling to the AT1 and AT2 cells (**Figure 11F**). TNFRSF12A^hi^ cells clustered within terminal bronchioles and exhibited enriched clonogenic distal lung organoid growth activity (103).

**Figure 11:**
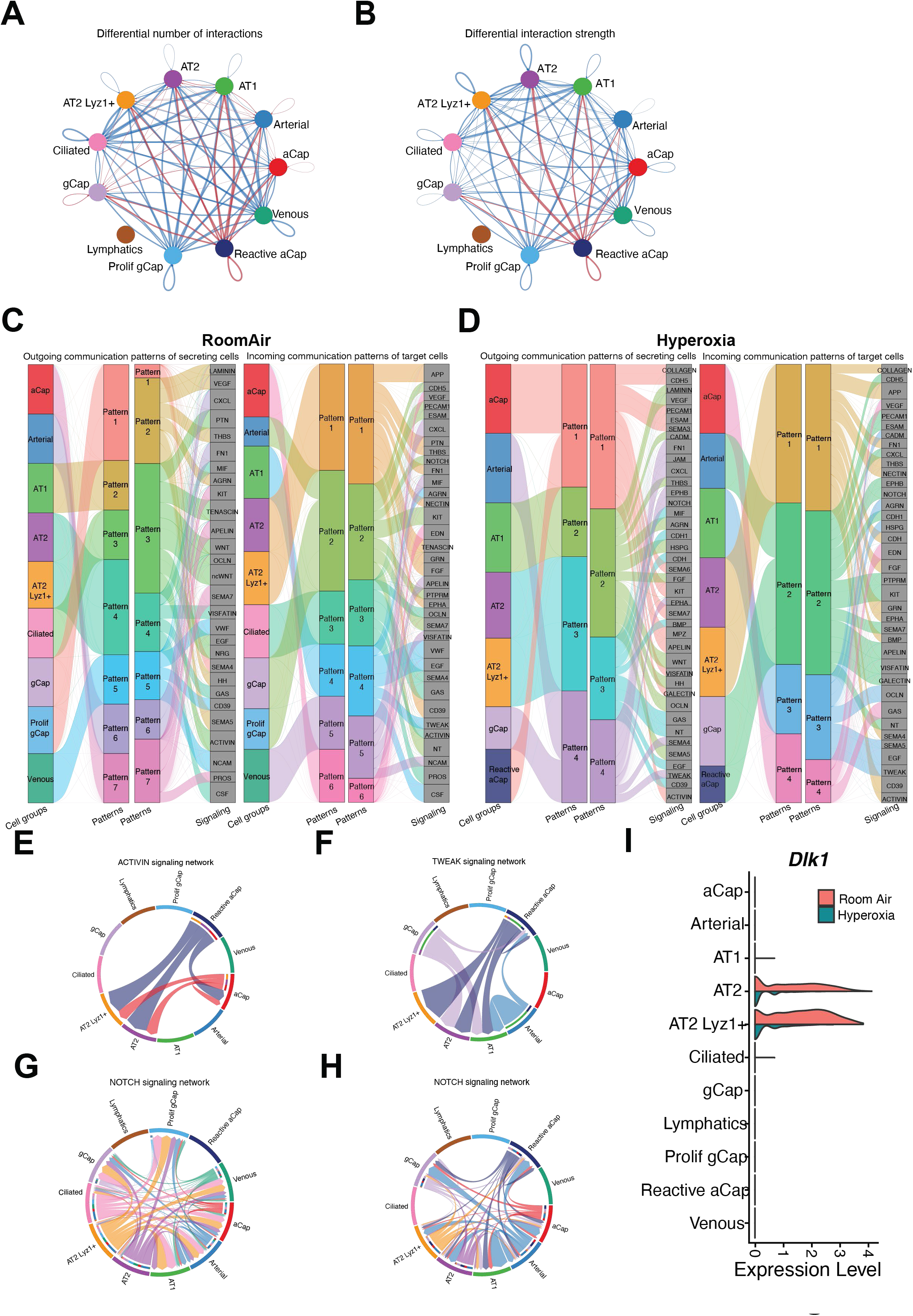
Neonatal hyperoxia decreases and alters ligand-receptor interactions between endothelial and epithelial cells. Circle plot showing the **A)** number and **B)** strength of differential interactions between epithelial and endothelial cell sub-populations in room air (blue) and hyperoxia (red). The inferred outgoing patterns of secreting cells and incoming communication patterns of target cells between the epithelial and endothelial cells, in **C)** room air and **D)** hyperoxia. The thickness of the flow indicates the contribution of the cell group or signaling pathway to each latent pattern. **E)** Chord diagram showing senders and targets for the activin signaling network under hyperoxia. The *activin* signaling network signals mainly from the reactive aCaPs to AT2 and AT2 Lyz1+ cells. **F)** Chord diagram showing senders and targets for the *Tweak* signaling network under hyperoxia. This was another inferred endothelial to epithelial communication with the ligand from the gCaPs, aCaP, and arterial endothelial cells and signaling to the AT1 and AT2 cells. Chord diagram showing senders and targets for the *Notch* signaling network under **G)** room air and **H)** hyperoxia. The epithelial to endothelial interaction in the Notch signaling pathway is decreased in hyperoxia. **I)** Violin plots showing expression of *Dlk1* (Notch ligand) in lung epithelial and endothelial cells in room air and hyperoxia.

Blockade of the Tweak receptor decreased lung fibrosis and mortality in a sepsis-induced acute lung injury model (85). The density of epithelial to endothelial interactions in the Notch signaling pathway is decreased in hyperoxia (**Figure 11 G-H**). One of the Notch ligands identified in the epithelial cells was *Dlk1* (Delta like non-canonical Notch ligand 1). *Dlk1* expression was high in AT2 cells in room air with a significant decrease in hyperoxia (**Figure 11I**). *Dlk1* regulated lung epithelial cell proliferation and differentiation during fetal lung development (104). *Dlk1* signaling was also necessary for timed inhibition of Notch signaling to allow AT2-AT1 differentiation (105). Interestingly, an inhibitory role on angiogenesis has been ascribed to *Dlk1* (*106*).

### Neonatal hyperoxia increases and alters ligand-receptor interactions between endothelial and immune cells

The number of inferred interactions is increased between the endothelial and immune cells with a decrease in the strength of interactions in hyperoxia compared to room air **(Supp Figure 5A).** aCaPs and reactive aCaPs showed the highest outgoing interaction strength, while dendritic cells, interstitial macrophages and classical monocytes the highest incoming interaction under normoxia and hyperoxia **(Supp Figure 5B).** Under exposure to hyperoxia, the predominant endothelial cell population serving as the signaling hub to immune cells were the reactive aCaPs **(Figure 12A).** The communication patterns between immune and endothelial cells in room air and hyperoxia are shown in **Supp Figure 5 C-D**. Under room air, four outgoing patterns are observed with all the immune cells except NK cells, B cells, neutrophils 2, and T cells forming the predominant secreting cell group (pattern #2). Under hyperoxia, the outgoing signaling patterns condensed to five main groups with all the immune cells forming the predominant secreting cells group (pattern #2). Among the endothelial cells, the aCaP/reactive aCaP (pattern #1) and the gCaP and arterial endothelium (pattern #5) formed distinct groups of secreting cells. There was no major VEGF signaling between endothelial and immune cells in room air (**Figure 12B**). Under hyperoxia, the neutrophil 3 population signals to the distal capillary endothelium with VEGF as the proposed ligand (**Figure 12C-D**). Neutrophils as a source of VEGF and mediating angiogenesis in pathological conditions has been reported previously (107–109). Another proposed ligand from basophils, interstitial macrophages and neutrophils 3 signaling to the distal lung endothelium upon exposure to hyperoxia was TNF-alpha (**Figure 12E-F**). TNF-alpha-induces expression of endothelial cell adhesion molecules such as ICAM1 (110, 111). It also modulates blood vessel remodeling in the setting of inflammation, playing a role in pathological angiogenesis (112, 113) . *Tnf* expression has been reported in lung basophils(95) and protective in injury models as well (114).

**Figure 12:**
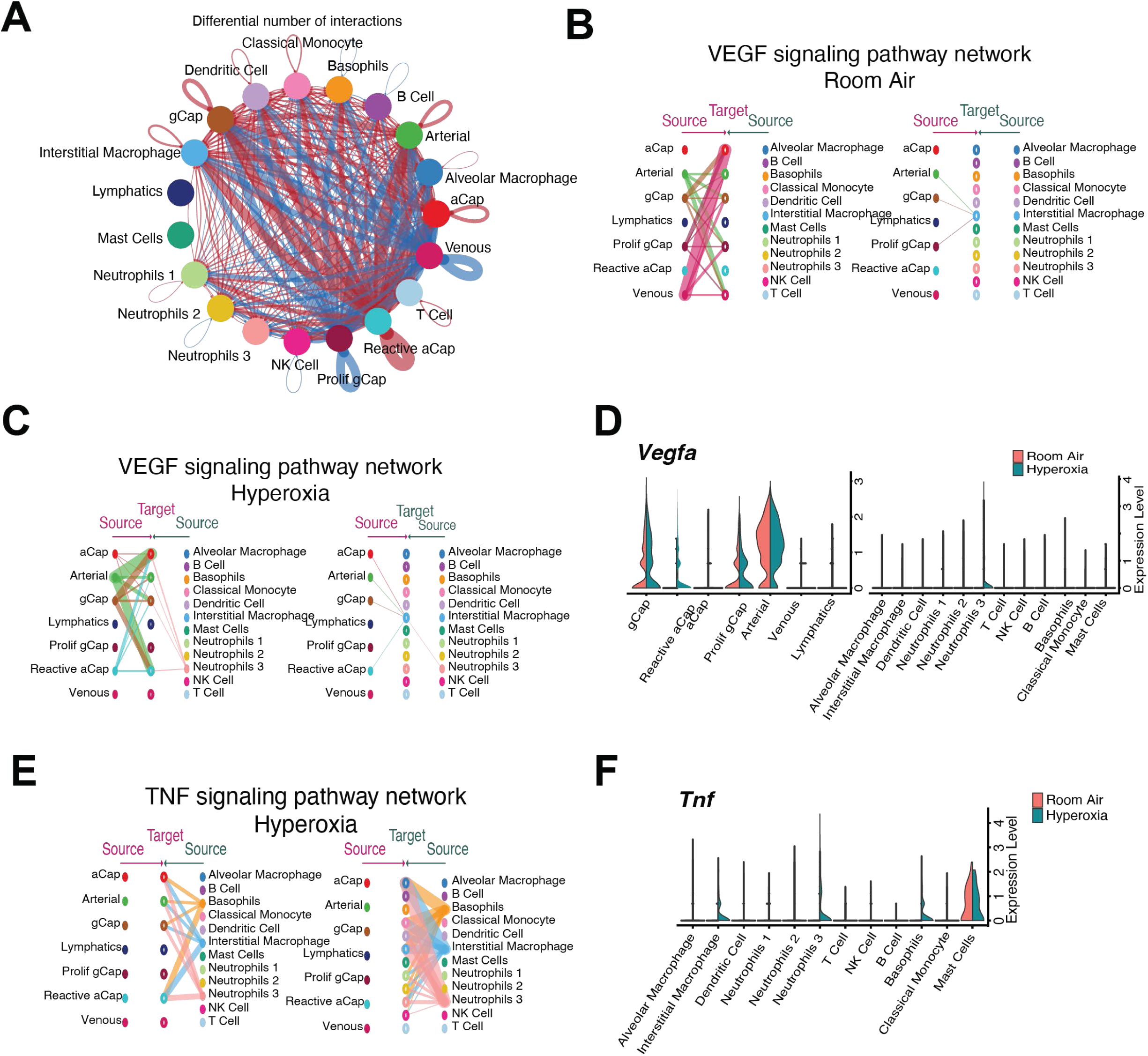
Neonatal hyperoxia increases and alters ligand-receptor interactions between endothelial and immune cells. **A)** Circle plot showing the differential number of interactions between different lung endothelial and immune cell sub-populations in room air (blue) and hyperoxia (red). **B)** Hierarchical plot showing the inferred intercellular communication network for *Vegf* signaling between the endothelial and immune cells in room air and **C)** hyperoxia. **D)** Violin plots showing expression of *Vegf* in lung endothelial and immune cells in room air and hyperoxia. **E)** Hierarchical plot showing the inferred intercellular communication network for *Tnf* signaling between the endothelial and immune cells in hyperoxia. This plot consists of two parts: Left portions highlights signaling from immune cells to endothelial cells. The right portion shows signaling within immune cells. Solid and open circles represent source and target, respectively. Circle sizes are proportional to the number of cells in each cell group and edge width represents the communication probability. Edge colors are consistent with the signaling source. **F)** Violin plots showing expression of *Tnf* in lung endothelial and immune cells in room air and hyperoxia.

## DISCUSSION

We present significant sex-specific differences in all lung cell sub-populations by sc-RNA seq of the PND7 murine lung at the early alveolar stage of lung development following hyperoxia exposure during the saccular stage of lung development. We and others have previously shown that the female lung is more resilient to neonatal hyperoxia exposure with better preservation of alveolarization and vascular development compared to similarly exposed neonatal male lung (17–19). Sex-specific differences by whole lung bulk RNA-Seq revealed some differences at PND7, which were increased during recovery at PND21 (20). Our objective was to delineate the sex-specific differences in the postnatal hyperoxia murine model at single-cell resolution to highlight the sexual dimorphism in different lung cell sub- populations. Sexual dimorphism was evident in all lung subpopulations but was striking among the lung immune cells. Finally, we identified that the specific intercellular communication networks and the ligand-receptor pairs that are impacted by neonatal hyperoxia exposure. Xia *et al.* have reported sex-specific differences in the lung cell sub- populations using different hyperoxia exposure model using 85% FiO2 for 14 days after birth (115).

Postnatal hyperoxia had a significant impact on the distal lung capillary endothelium compared to the arterial/venous endothelium with a remarkable change in the cell state of aCaPs and gCaPs. We show upregulation of *Inhba* in the reactive aCaP capillary endothelial cells upon exposure to hyperoxia. Inhibins belong to the TGF-beta family and is comprised of a functional heterodimer with an alpha- and a beta-subunit. Inhibin promotes angiogenesis through TGF-beta receptors (116) and increases vascular permeability through internalization of VE-cadherin (117). Under hypoxic conditions, Inhibin expression is modulated *HIF-1a* and *HIF-2a* (118). However, an abundance of *Inhba* impairs endothelial cells function by reducing BMPPR2 levels in endothelial cells (98) . The reactive aCaPs seem to be a transient endothelial cell population arising from the gCaP cells and leading to the aCaP cells with markers of both aCaP and gCaP cells. Whether this endothelial cell subpopulation is seen in later time points during recovery and repair and in the human disease in preterm neonates needs to be elucidated.

We identified three distinct neutrophil subpopulations in our data set and highlight the existence of the polymorphonuclear neutrophil myeloid derived suppressor cells (PMN-MDSC) in the neonatal lung and the emergence of activated PMN-MDSCs upon exposure to hyperoxia, highlighting the heterogeneity in lung neutrophils at baseline, after exposure to hyperoxia and differences between the male and female lung. MDSCs play a crucial role in immunosuppression in cancer, by production of reactive oxygen species, immunosuppressive cytokines and adversely impacting T-cell proliferation (119) . Several mediators including M-CSF, which was upregulated in the hyperoxia exposed lung at PND7 can promote the development of MDSCs. We show three distinct populations of neutrophils among the room air and hyperoxia-exposed lung with neutrophils 1 (*Ngp+ Camp+ and Lcn2+)*, neutrophils 2, and the neutrophils 3 which were enriched for genes related to cell activation and inflammation (*Ccl4, Ccl3, Cxcl3, Il1b, Nfkbia, Mif, Klf6, Atf3*). Neutrophils 3 emerge in the hyperoxia exposed lung. Velocity analysis showed that the neutrophils 3 arise from neutrophils 2. The neutrophils 3 exhibit a transcriptional state reminiscent of the activated PMN-derived MDSCs (26). Neutrophils 3 showed the greatest number of differentially expressed genes in response to hyperoxia and also produced VEGF and signal to the lung endothelium.

Significantly, we highlight the sex-specific differences in gene expression in different lung cell sub-populations in the epithelial, endothelial, and immune cells. Differences in the secretome and transcriptome of human umbilical venous endothelial cells has been described (120) and so is the response to hyperoxia in human pulmonary microvascular endothelial cells (121). The immune cells stand out in the degree of sexual dimorphism in the hyperoxia exposed neonatal lung. The sexual dimorphism in both innate and adaptive immune response in many diseases and organs is well known (122, 123). Interestingly, many immune-related genes and regulatory elements that play a role in both the innate and adaptive immune responses are located on the X-chromosome (124). Apart from the highlighted differentially expressed genes that are distinct between the male and the female lung, several sex-specific biological pathways, that were common between different lung cell sub-populations were discovered. Of note, apoptosis, TNF-alpha signaling pathway, p53 pathway, inflammatory response, interferon-gamma response and cytokine mediated signaling was downregulated in females. Electron transport chain, inflammatory response, apoptosis and TNF-alpha signaling was upregulated in males. Differences between male and female cells in changes in mitochondrial respiration upon exposure to various stressors and the sex-dependent impacts have been reported in many previous studies (125–128). Male-specific upregulation of TNF-alpha was described in response to other injuries and other organs (129–132) in newborns and adults. The differences between the male and female-specific biological pathways corresponds with the phenotype of greater lung injury with decreased alveolarization and vascular development in the hyperoxia-exposed male lung compared to the female (18).

We highlighted cell-cell communications between the major lung cellular sub- populations and have highlighted novel autocrine and paracrine acting ligands and ligand receptor pairs. This resource will be made available to scientific community at https://www.lingappanlab.com/resources. The aCaPs and reactive aCaPs function as the main senders among the lung endothelial cells, while the AT1 cells serve that role among epithelial cells. Interstitial macrophages and classical monocytes serve as the signaling hubs among the lung immune cells. The number of intercellular communications as well as the strength of communication is generally decreased upon exposure to hyperoxia except among endothelial cells and immune cells, where the number of interactions is increased. We highlight several known and well-established signaling networks such as *Notch, TGF-beta, Bmpr2 and Tnf-alpha* signaling networks, but at the same time were also able to reveal novel ligand and ligand-receptor interactions such as the *Tweak* and *Ptprm* signaling networks.

We recognize the limitations of the present study with the use of pooled lung cells from three biological replicates. The lung dissociation methods may have biased our yield with increased isolation of certain lung dell subpopulations, while decreasing the yield of others. Future studies would investigate the sex-specific transcriptional state in the lung at prenatal and at P1 (at baseline) before the onset of injury, that may predispose the male and female lung to different patterns of lung injury postnatally. We had previously identified that the sex-specific differences in gene expression increase at late alveolar stage of lung development (PND21) following hyperoxia exposure during the saccular stage of lung development. Cell states during recovery and repair after early injury at single-cell resolution need to be elucidated. Sex-specific differences may be mediated through sex hormones or through genes on the X-chromosome having differential expression in the male and female lung. Exploring the basis behind sex-specific differences will be crucial to explain the female sex resilience in human BPD and will suggest new therapeutic modalities and guide the right therapy to the right patient.

## Acknowledgements

This work was supported in part by grants from National Institutes of Health [R01-HL144775; R01-HL146395 and R21-HD100862 to KL]. We acknowledge Dr. Lukas Simon for his help with initial approaches for data analysis. We also acknowledge Dominique Armstrong from helping with the single-cell isolation.

## Supplemental Data

### Supplemental Tables

**Supplemental Table 1:** The number of cells sequenced in each group and the highly expressed genes in each cluster

**Supplemental Table 2:** Differentially expressed genes in all endothelial cell sub- populations by experimental condition and sex.

**Supplemental Table 3:** Differentially expressed genes in all epithelial cell sub-populations by experimental condition and sex.

**Supplemental Table 4:** Differentially expressed genes in all immune cell sub-populations by experimental condition and sex.

Supplemental Figures:

**Supplemental Figure 1: A)** In situ hybridization and violin plots showing the increased expression of *Peg 3* in aCaP (*Ednrb+*) lung endothelial cells in response to hyperoxia. **B)** In situ hybridization and violin plots showing the increased expression of *Ly6a* in gCaP (*Aplnr+*) lung endothelial cells in response to hyperoxia. Pathway analysis showing pathways enriched among upregulated (blue) or downregulated (yellow) differentially expressed genes in **C)** aCaP/reactive aCaP and **D)** gCaP cells.

**Supplemental Figure 2:** Pathway analysis showing pathways enriched among upregulated (blue) or downregulated (yellow) differentially expressed genes in **A)** interstitial macrophages and **B)** Neutrophils 3. **C)** Volcano plots showing up- and down- regulated genes in female classical monocytes. **D)** Sex-specific enriched biological pathways (blue: male, pink: female) in lung classical monocytes upon exposure to hyperoxia

**Supplemental Figure 3: A)** The number of interactions and interaction strength between lung endothelial cells in room air and hyperoxia. **B)** The number of interactions and interaction strength between lung epithelial cells in room air and hyperoxia. **C)** To visualize the dominant senders (sources) and receivers (targets) in room air and hyperoxia, in a 2D space, we generated a scatter plot.

**Supplemental Figure 4: A)** The number of interactions and interaction strength between lung immune cells in room air and hyperoxia. **B)** To visualize the dominant senders (sources) and receivers (targets) in room air and hyperoxia, in a 2D space, we generated a scatter plot.

**C)** The number of interactions and interaction strength between lung endothelial and epithelial cells in room air and hyperoxia. **D)** To visualize the dominant senders (sources) and receivers (targets) in room air and hyperoxia, in a 2D space, we generated a scatter plot.

**Supplemental Figure5: A)** The number of interactions and interaction strength between lung endothelial and epithelial cells in room air and hyperoxia. **B)** To visualize the dominant senders (sources) and receivers (targets) in room air and hyperoxia, in a 2D space, we generated a scatter plot. The inferred outgoing patterns of secreting cells and incoming communication patterns of target cells between the epithelial and endothelial cells, which shows the correspondence between the inferred latent patterns and cell groups, as well as signaling pathways in **C)** room air and **D)** hyperoxia. The thickness of the flow indicates the contribution of the cell group or signaling pathway to each latent pattern.

## MATERIALS AND METHODS

### Mice

All animal experiments were performed under an approved protocol by the IACUC at the Children’s Hospital of Philadelphia. Timed pregnant C57BL/6N WT mice were obtained from Charles River Laboratories (Wilmington). The sex in neonatal mouse pups was determined by both the anogenital distance and pigmentation in the anogenital region method. In neonatal male mice, a pigmented spot on the scrotum is visible to the naked eye from postnatal day (PND) 1, whereas female pups lack visible pigmentation in the anogenital region. This was also verified by the expression of Y-chromosome transcript abundance in male samples.

### Mouse model of BPD

An arrest of alveolarization was induced in mouse pups by exposure to hyperoxia (95% O2), as described previously(23). Mouse pups from multiple litters were pooled before being randomly and equally redistributed to two groups, one group exposed to room air (21% O2) and the other group exposed to hyperoxia (95% O2), within 12 h of birth for 5 days. The dams were rotated between air- and hyperoxia-exposed litters every 24h.

### Lung Isolation for Single Cell Sequencing

Mice were euthanized on post-natal day (PND) 7 with i.p. pentobarbital. The right ventricle was perfused with ice cold PBS and the lungs were harvested quickly and homogenized and dissociated using the Miltenyi MACS lung dissociation kit (Miltenyi Biotec, cat # 130-095-927), using the GentleMacs program m-Lung-1 on the AUTO MaCs. The lungs were incubated for 20min at 37 ◦C under continuous rotation and then run on the gentleMACS m-Lung-2 program on the gentleMACS (Miltenyi Biotec, Cat #130-095-937). This was followed by brief centrifugation and the passage of cells through the 70-uM filter, washed, centrifuged again and the pellet was treated with ACK lysis buffer, centrifuged, and resuspended in sorting buffer and passed through an 40uM filter. Live cells were sorted and subjected to single-cell RNA sequencing as described below.

### scRNA-Seq Library Preparation and Sequencing

Single cell Gene Expression Library was prepared according to Chromium Single Cell Gene Expression 3v3.1 kit (10x Genomics). In Brief, single cells, reverse transcription (RT) reagents, Gel Beads containing barcoded oligonucleotides, and oil were loaded on a Chromium controller (10x Genomics) to generate single cell GEMS (Gel Beads-In-Emulsions) where full length cDNA was synthesized and barcoded for each single cell. Subsequently the GEMS are broken and cDNA from each single cell are pooled. Following cleanup using Dynabeads MyOne Silane Beads, cDNA is amplified by PCR. The amplified product is fragmented to optimal size before end-repair, A-tailing, and adaptor ligation. Final library was generated by amplification. 10X Genomics Cell Ranger [version] ‘cellranger count’ was used to perform alignment, filtering, barcode counting, and UMI counting. The raw data has been uploaded to NCBI GEO; GSE211356.

### Sequencing, data processing, quality control, integration and cluster annotation

For each sample, a Seurat object was created using Cellranger output matrices using Read10X method. SCTransformV2 and Get clusters (RunPCA, RunUMAP, RunTSNE, FindNeighbors, FindClusters) was run. Then we filtered using SoupX version 1.6.1 (133)with a contamination factor of 5%. We used PercentageFeatureSet to get percent of mitochondrial genes in each cell. Resulting cells were filtered using >= 500 molecular identifiers (nUMI >= 500), removing upper number of UMI outliers (nUMI <= outlier_threshold_nUMI), >250 detectable genes (nGene >= 250), removing upper number of gene outliers (nGene <= outlier_threshold_nGene), log10GenesPerUMI > 0.80, < 5% of transcripts coming from mitochondrial genes (mitoRatio < 0.05). Genes were filtered, only keeping those genes expressed in more than 10 cells. Genes ’*Gm42418*’, ’*S100a8*’, ’*S100a9*’ (134, 135) were removed from Seurat object. We ran SCTransformV2 and Get clusters, resolution 0.6 was chosen. Doublets were detected and removed using DoubletFinder version 2.0.3 (136). paramSweep_v3, summarizeSweep, find.pK, doubletFinder_v3. Get clusters ran again. We aggregated the samples using SelectIntegrationFeatures with Number of features to return set to 3000, PrepSCTIntegration, FindIntegrationAnchors with sample 1 as reference (Female Hyperoxia), and IntegrateData using SCT as normalization method. Ran Get clusters again and chose resolution 0.6. Ran PrepSCTFindMarkers with default parameters and used the FindMarkers function for each cluster with parameters set to Only return positive markers using wilcoxon test and SCT assay. Differential expression between conditions were found using Seurat’s FindMarkers function. Genes detected in 25% of cells with an FDR (adjusted p-value) value of < 0.05 and a log2fold change of > 0.3 were reported as significant in each comparison. Significant genes were then processed using enrichR (version 3.0) (137) along fgsea (version 1.22.0) (138) to find enriched biological pathways present. RNA velocity estimation was done using cellranger output files, then sorted with samtools (1.14), then used velocyto (0.17)(139), then scVelo (0.2.4)(140). Cell-cell communication was analyzed using Cellchat (version 1.4.0) (83) on room air and hyperoxia samples separately with the software’s default parameters.

### Other Software Used

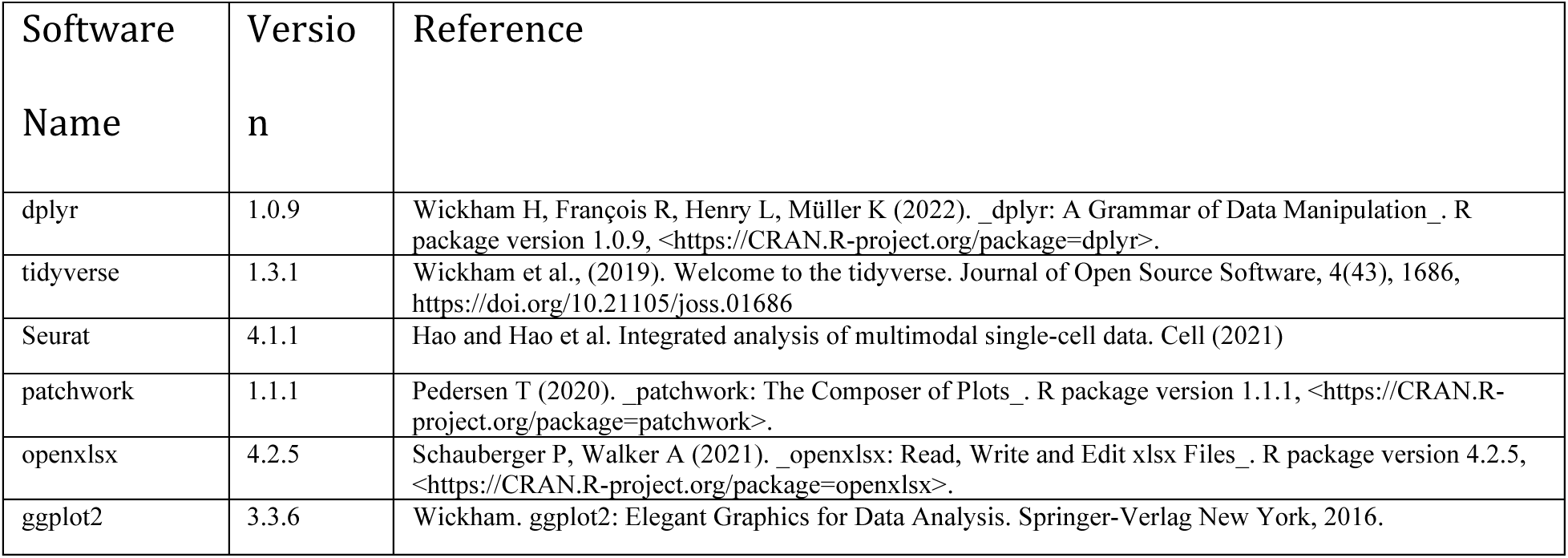
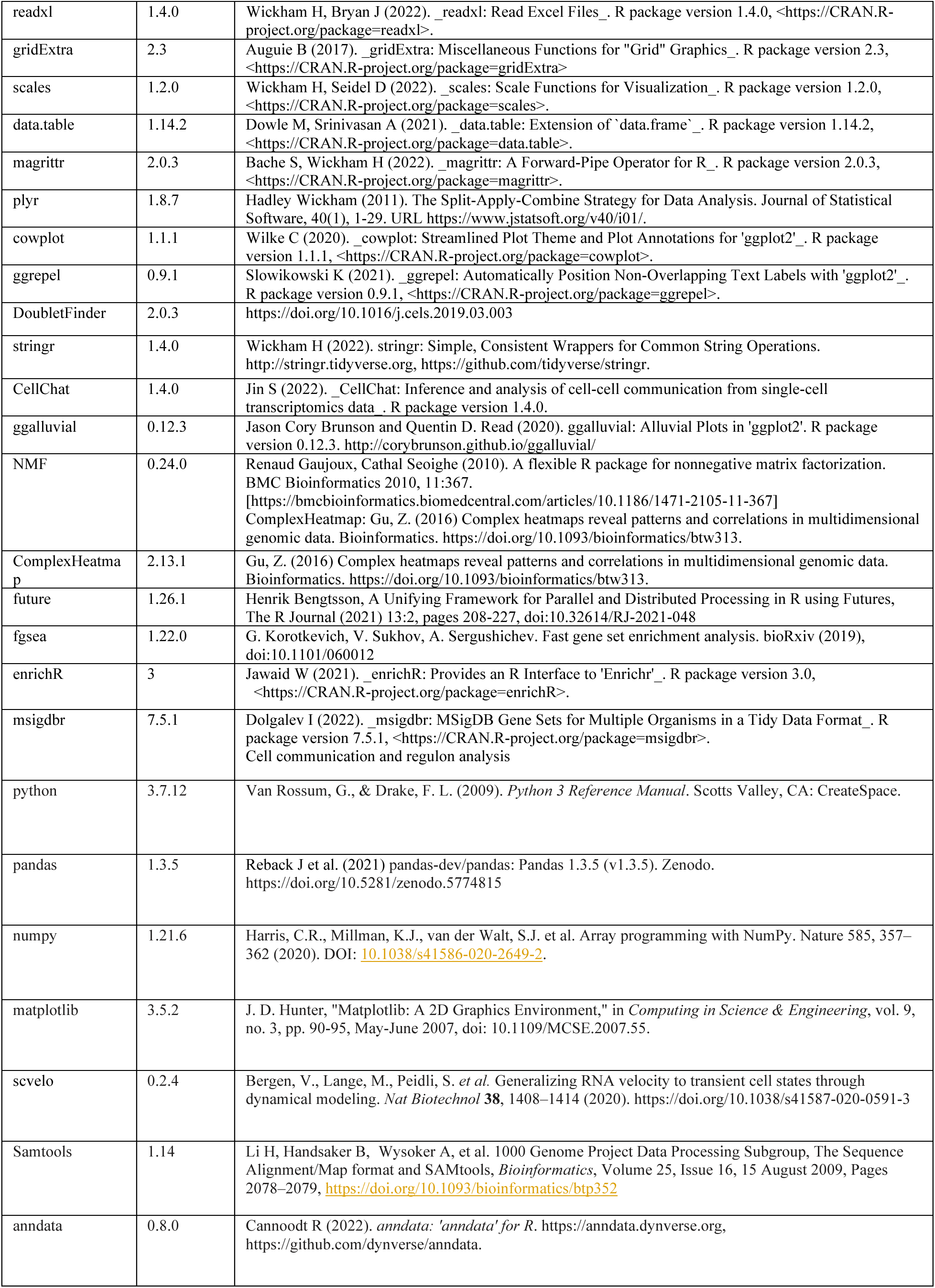
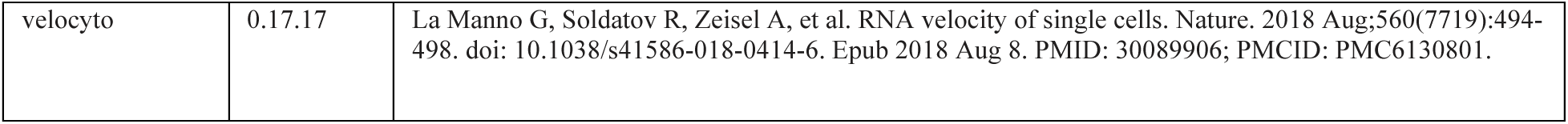

### In Situ Hybridization

*In situ* hybridization was performed using the RNAscope Multiplex Fluorescent Reagent Kit v2 (Advanced Cell Diagnostics, Hayward, CA, USA). Samples were treated with *Pecam (*316721)*, Inhba (*455871-C4)*, Ednrb (*473801-C2)*, Peg3 (*492581-C3)*, Aplnr (*436171)*, and Ly6A (*427571-C2) probes (Advanced Cell Diagnostics, Hayward, CA, USA) for 2 hours at 4°C. For each round, samples were run alongside a positive control slide treated with housekeeping genes *Polr2A, Ppib*, *Ubc, and Hprt (*321811) and a negative control slide which was treated with dapB (a bacteria gene) (321831). Nuclei were counterstained with DAPI and autofluorescence was reduced using TrueBlack Plus Lipofuscin Autofluoresne Quencer 40X in DMSO (Biotum: 23014) for 30 minutes then washed in PBS. Slides were mounted in EverBrite TrueBlack® Hardset Mounting Medium (Biotium: 23017). The following day, slides were imaged using Leica Bmi8 Thunder Imager microscope.

## Notes

### Competing Interest Statement

The authors have declared no competing interest.

