## Supplemental Figures for "Remarkable sex-specific differences at Single-Cell Resolution in Neonatal Hyperoxic Lung Injury"

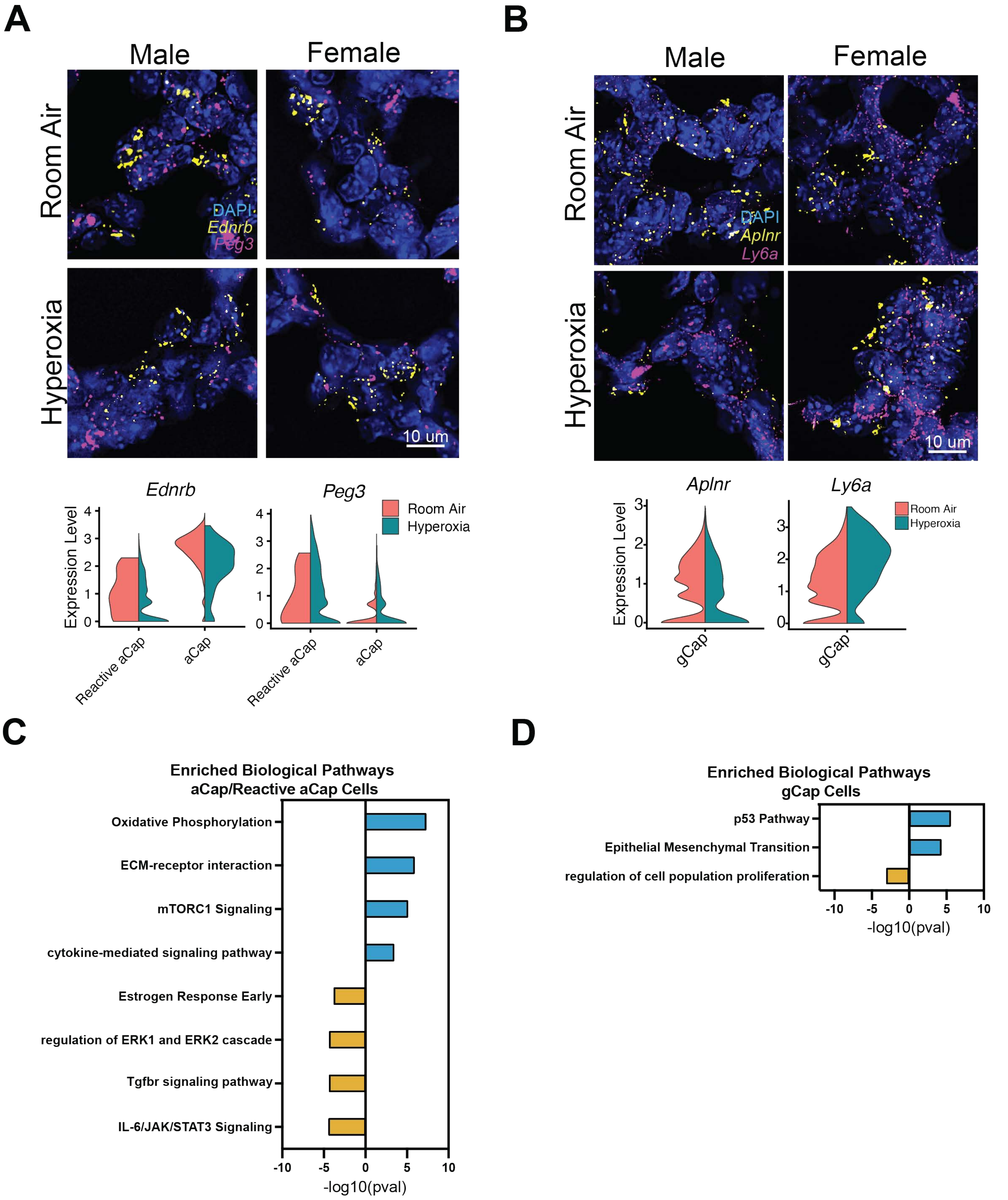


**Supplemental FIgure 1**

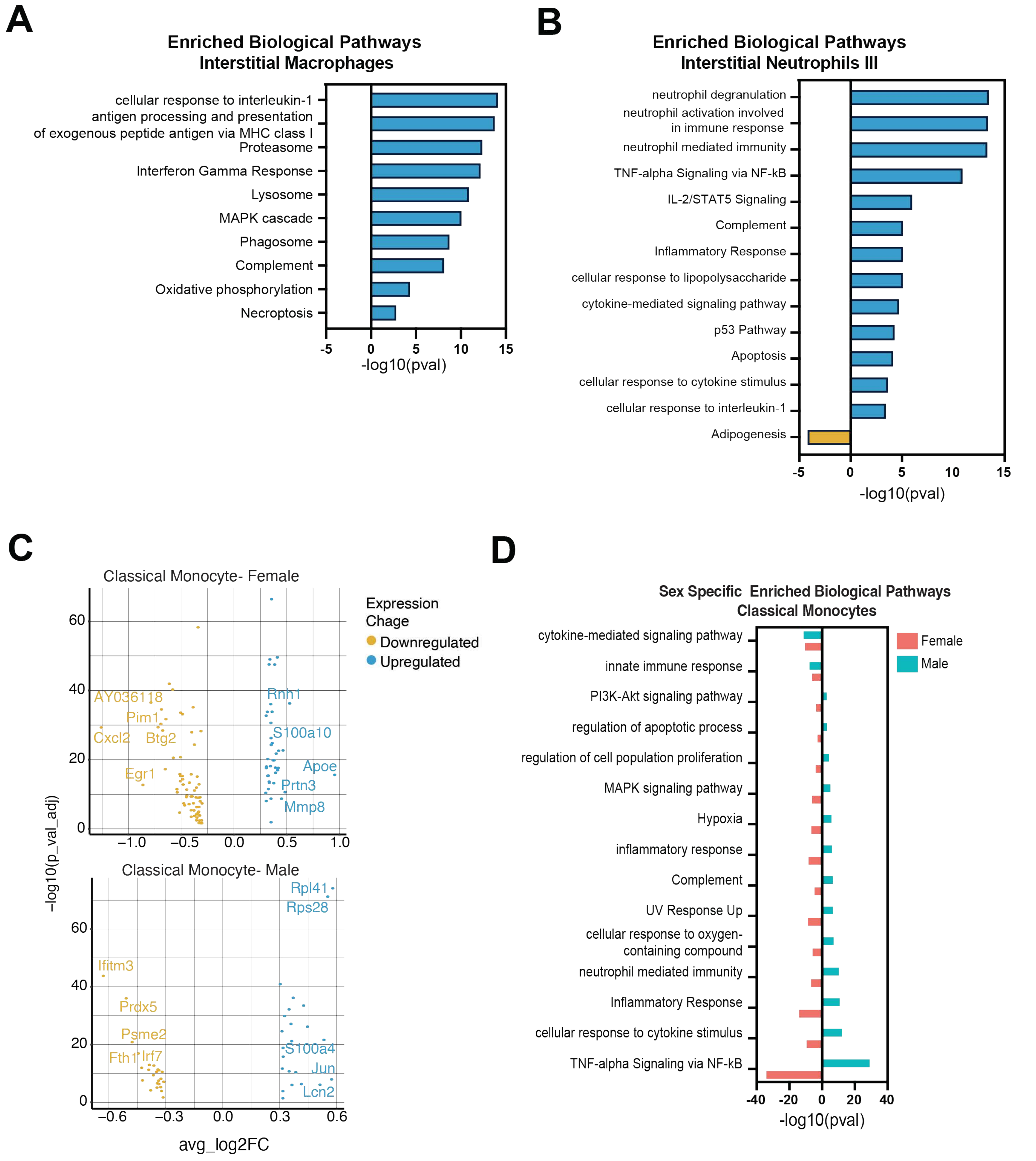


Supplemental Figure 2

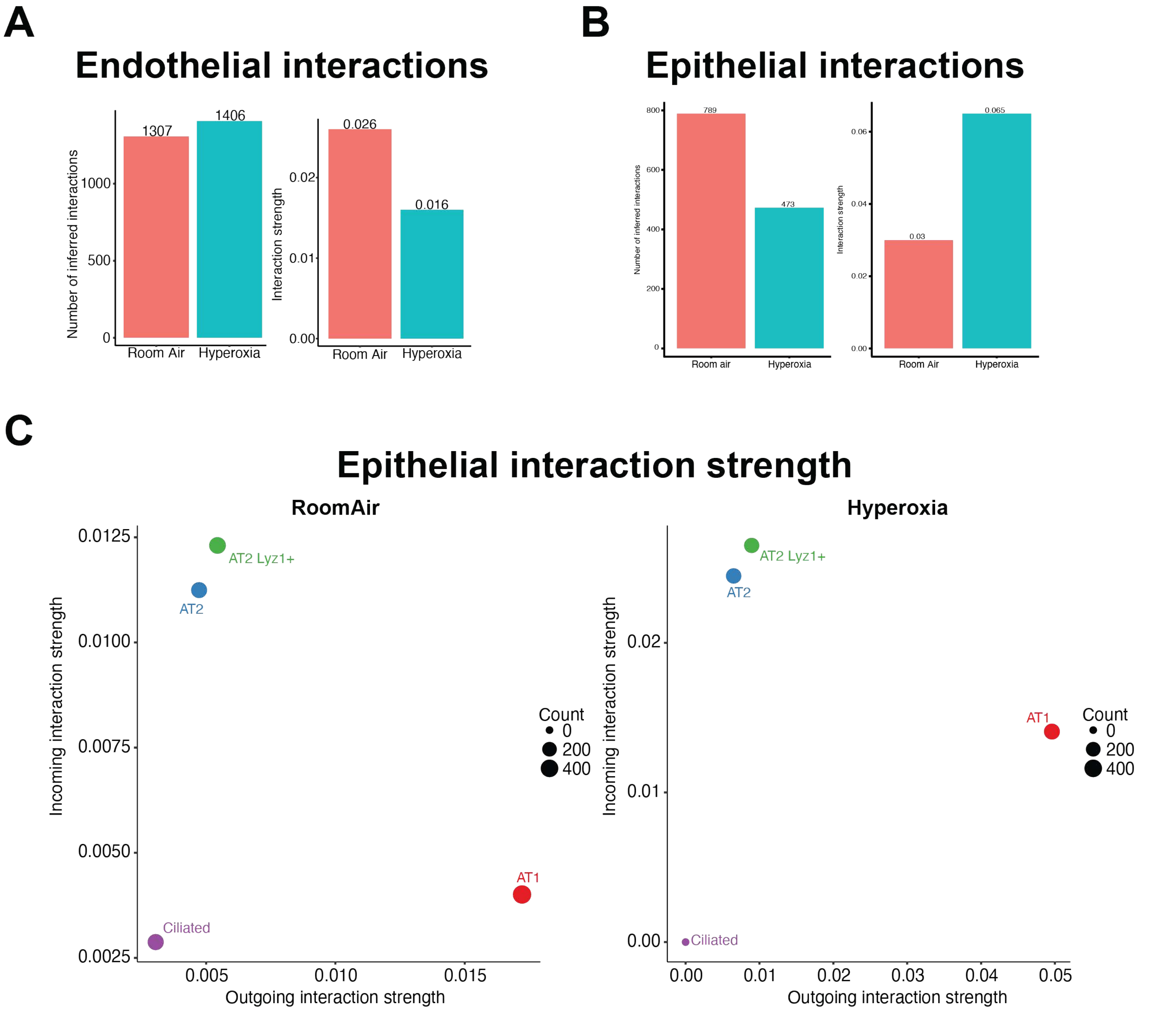


**Supplemental Figure 3**

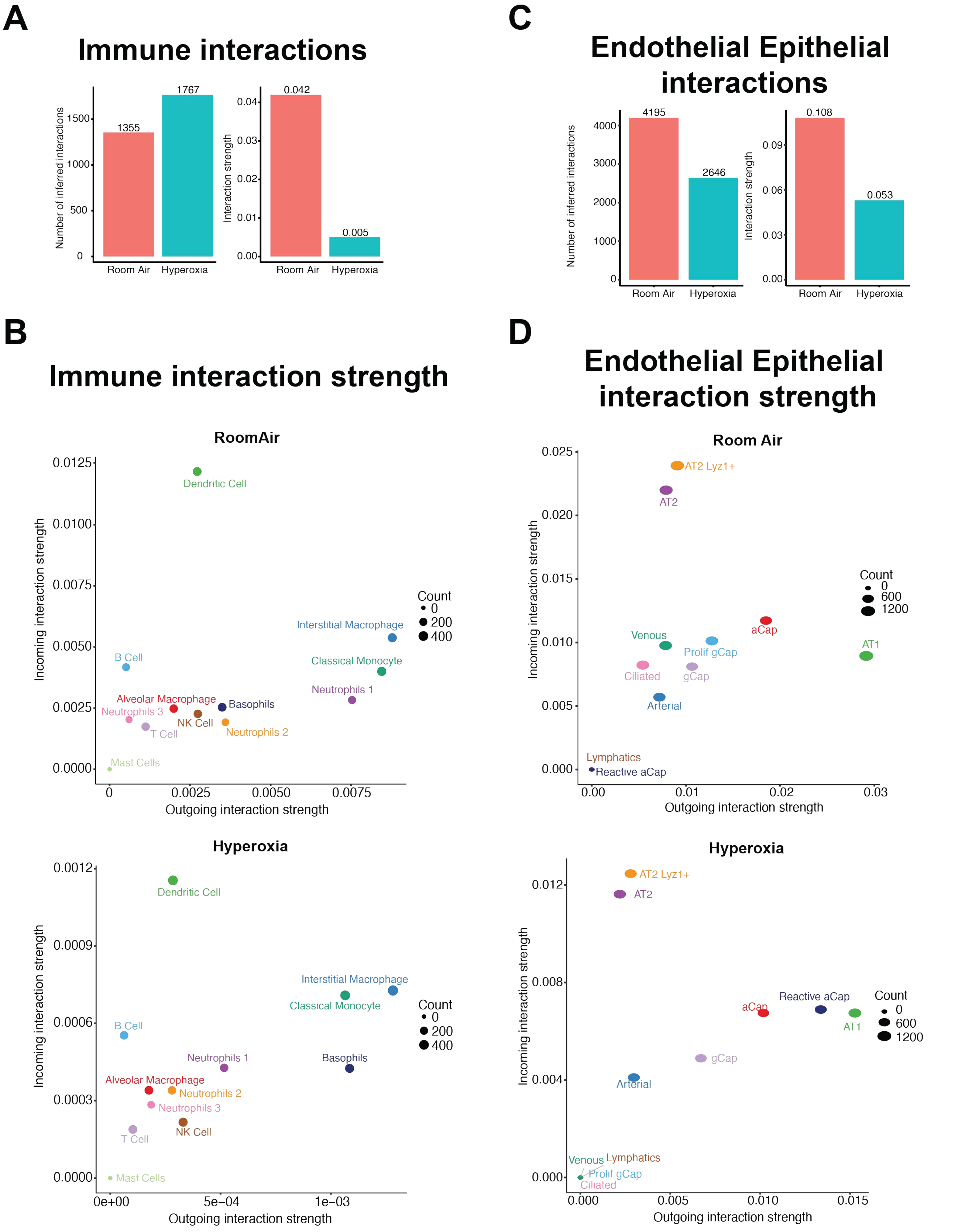


**Supplemental Figure 4**

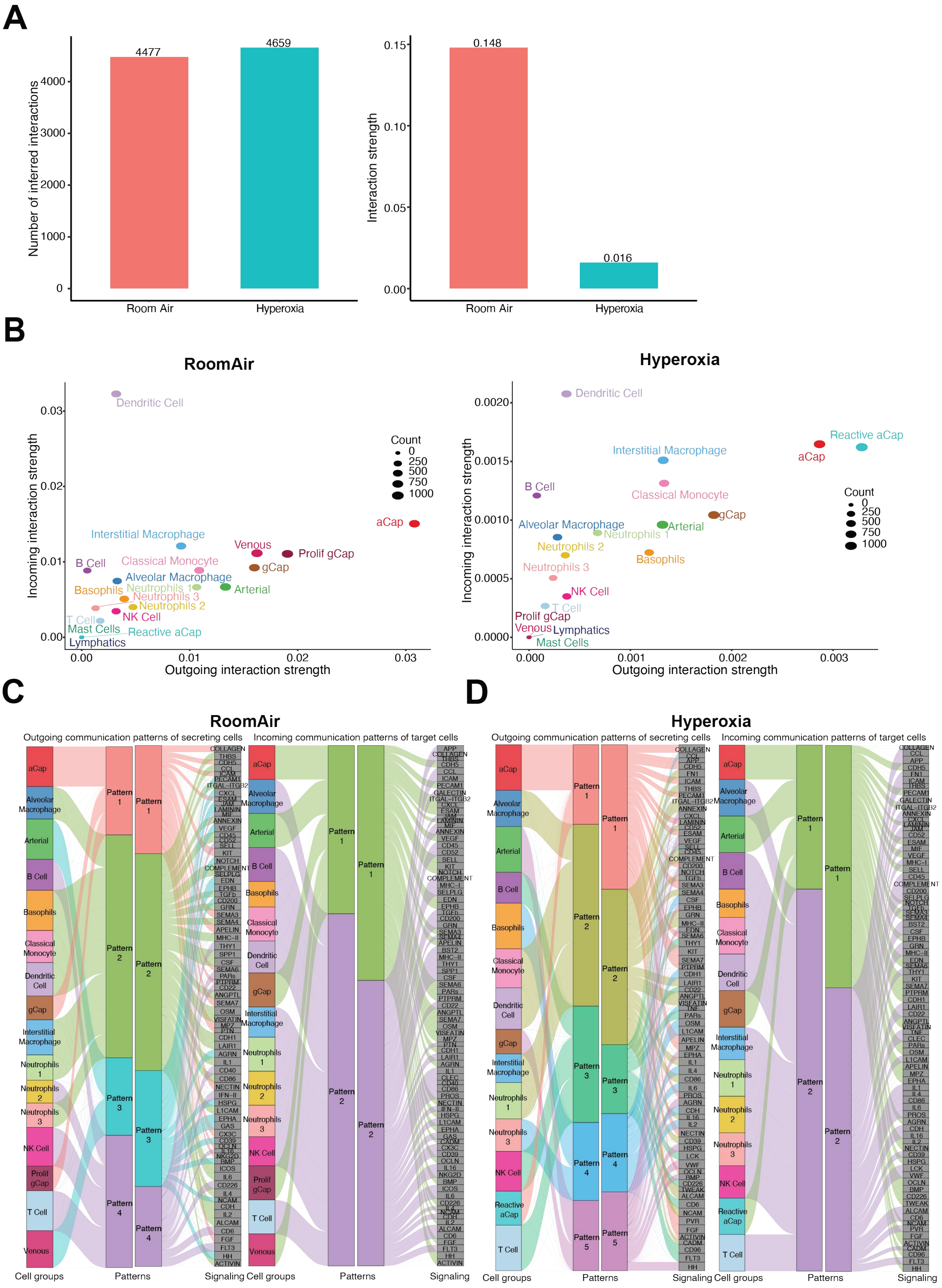


**Supplemental Figure 5**
